# Global Genetic Cartography of Urban Metagenomes and Anti-Microbial Resistance

**DOI:** 10.1101/724526

**Authors:** David Danko, Daniela Bezdan, Ebrahim Afshinnekoo, Sofia Ahsanuddin, Chandrima Bhattacharya, Daniel J Butler, Kern Rei Chng, Daisy Donnellan, Jochen Hecht, Katelyn Jackson, Katerina Kuchin, Mikhail Karasikov, Abigail Lyons, Lauren Mak, Dmitry Meleshko, Harun Mustafa, Beth Mutai, Russell Y Neches, Amanda Ng, Olga Nikolayeva, Tatyana Nikolayeva, Eileen Png, Krista Ryon, Jorge L Sanchez, Heba Shaaban, Maria A Sierra, Dominique Thomas, Ben Young, Omar O. Abudayyeh, Josue Alicea, Malay Bhattacharyya, Ran Blekhman, Eduardo Castro-Nallar, Ana M Cañas, Aspassia D Chatziefthimiou, Robert W Crawford, Francesca De Filippis, Youping Deng, Christelle Desnues, Emmanuel Dias-Neto, Marius Dybwad, Eran Elhaik, Danilo Ercolini, Alina Frolova, Dennis Gankin, Jonathan S. Gootenberg, Alexandra B Graf, David C Green, Iman Hajirasouliha, Mark Hernandez, Gregorio Iraola, Soojin Jang, Andre Kahles, Frank J Kelly, Kaymisha Knights, Nikos C Kyrpides, Paweł P Łabaj, Patrick K H Lee, Marcus H Y Leung, Per Ljungdahl, Gabriella Mason-Buck, Ken McGrath, Cem Meydan, Emmanuel F Mongodin, Milton Ozorio Moraes, Niranjan Nagarajan, Marina Nieto-Caballero, Houtan Noushmehr, Manuela Oliveira, Stephan Ossowski, Olayinka O Osuolale, Orhan Özcan, David Paez-Espino, Nicolas Rascovan, Hugues Richard, Gunnar Rätsch, Lynn M Schriml, Torsten Semmler, Osman U Sezerman, Leming Shi, Tieliu Shi, Le Huu Song, Haruo Suzuki, Denise Syndercombe Court, Scott W Tighe, Xinzhao Tong, Klas I Udekwu, Juan A Ugalde, Brandon Valentine, Dimitar I Vassilev, Elena Vayndorf, Thirumalaisamy P Velavan, Jun Wu, María M Zambrano, Jifeng Zhu, Sibo Zhu, Christopher E Mason, The International MetaSUB Consortium

## Abstract

We have created a global atlas of 4,728 metagenomic samples from mass-transit systems in 60 cities across 3 years. This is the first systematic, worldwide study cataloging the urban microbial ecosystem. We identify taxonomically-defined microorganisms collected across three years. This atlas provides an annotated, geospatial profile of microbial strains, functional characteristics AMR markers, and novel genetic elements, including 10,928 viral, 1302 bacteria, and 2 archaea novel species. We identify 4,424 species of urban microorganisms and a consistent “core” of 31 species found in nearly all samples that is largely distinct from any human commensal microbiome. Profiles of AMR genes show geographic variation in type and density. Together, these results constitute a high-resolution, global metagenomic atlas, which enables the discovery of new genetic components, highlights potential forensic applications, and provides an essential first draft of the global AMR burden of the world’s cities.

## 1 Introduction

The high-density urban environment has historically been home to only a fraction of all people, with the majority living in rural areas or small villages. In the last two decades, the situation has reversed; 55% of the world’s population now lives in urban areas (Ritchie and Roser, 2020; United Nations, 2018). Since the introduction of germ theory and John Snow’s work on cholera, it has been clear that people in cities interact with microbes in ways that can be markedly different than in rural areas (Neiderud, 2015). Microbes in the built environment have been implicated as a possible source of contagion (Cooley et al., 1998) and certain syndromes, like allergies, are associated with increasing urbanization (Nicolaou et al., 2005). It is now apparent that cities in general have an impact on human health though the mechanisms of this impact are broadly variable and often little understood. Indeed, our understanding of microbial dynamics in the urban environment outside of pandemics has only begun (Gilbert and Stephens, 2018).

Technological advances in next-generation sequencing (NGS) and metagenomics have created an unprecedented opportunity for rapid, global studies of microorganisms and their hosts, providing researchers, clinicians, and policymakers with a more comprehensive view of the functional dynamics of microorganisms in a city. NGS facilitates culture-independent sampling of the microorganisms in an area with the potential for both taxonomic and functional annotation; this is particularly important for surveillance of microorganisms as they acquire antimicrobial resistance (AMR) (Fresia et al., 2019). Metagenomic methods enable nearly real-time monitoring of organisms, AMR genes, and pathogens as they emerge within a given geographical location, and have the potential to reveal hidden microbial reservoirs and detect microbial transmission routes as they spread around the world (Zhu et al., 2017). There are several different drivers and sources for AMR; including agriculture, farming, and livestock in rural and suburban areas, household and industrial sewage, usage of antimicrobials, hard metals, and biocides, as well as human and animal waste, all these factors contribute to the complexity of AMR transmission (Allen et al., 2009; Martínez, 2008; Singer et al., 2016; Thanner et al., 2016; Venter et al., 2017). A molecular map of urban environments will enable significant new research on the impact of urban microbiomes on human health.

The United Nations projects that by 2050, over two-thirds of the world’s population will live in urban areas (Ritchie and Roser, 2020). Consequently, urban transit systems – including subways and buses – are a daily contact interface for billions of people who live in cities. Notably, urban travelers bring their commensal microorganisms with them as they travel and come into contact with organisms and mobile elements present in the environment, including AMR markers. The study of the urban microbiome and the microbiome of the built environment spans several different projects and initiatives including work focused on transit systems (Afshinnekoo et al., 2015; Hsu et al., 2016; Kang et al., 2018; Leung et al., 2014; MetaSUB International Consortium. Mason et al., 2016), hospitals (Brooks et al., 2017; Lax et al., 2017), soil (Hoch et al., 2019; Joyner et al., 2019), and sewage (Fresia et al., 2019; Maritz et al., 2019), among others. However, these efforts for the most part have only been profiled with comprehensive metagenomic methods in a few selected cities on a limited number of occasions. This leaves a gap in scientific knowledge about a microbial ecosystem, with which the global human population readily interacts. Human commensal microbiomes have been found to vary widely based on culture, and thus the geography and geographically constrained studies may to miss key differences (Brito et al., 2016). Moreover, data on urban microbes and AMR genes are urgently needed in developing nations, where antimicrobial drug consumption is expected to rise by 67% by 2030 (United Nations, 2016; Van Boeckel et al., 2015), both from changes in consumer demand for livestock products and an expanding use of antimicrobials – both of which can alter AMR profiles of these cities.

The International Metagenomics and Metadesign of Subways and Urban Biomes (MetaSUB) Consortium was launched in 2015 to address this gap in knowledge on the density, types, and dynamics of urban metagenomes and AMR profiles. Since then, we have developed standardized collection and sequencing protocols to process 4,728 samples across 60 cities worldwide (Table S1). Sampling took place at three major time points: a pilot study in 2015-16 and two global city sampling days (gCSD, June 21st) in 2016 and 2017. Each sample was sequenced with 5-7M 125bp paired-end reads using Illumina NGS sequencers (see Methods). To deal with the challenging analysis of our large dataset, we generated an open-source analysis pipeline (MetaSUB Core Analysis Pipeline, CAP), which includes a comprehensive set of state-of-the-art, peer-reviewed, metagenomic tools for taxonomic identification, *k*-mer analysis, AMR gene prediction, functional profiling, de novo assembly, annotation of particular microbial species, and geospatial mapping.

To our knowledge this study represents the first and largest global metagenomic study of urban microbiomes – with a focus on transit systems – that reveals a consistent “core” urban microbiome across all cities, as well as distinct geographic variation that may reflect epidemiological variation and that enables a new forensic, source-tracking capabilities. More importantly, our data demonstrate that a significant fraction of the urban microbiome remains to be characterized. Though 1,000 samples are sufficient to discover roughly 80% of the observed taxa and AMR markers, we continued to observe taxa and genes at an ongoing discovery rate of approximately one new species (previously non-observed) and one new AMR marker for every 10 samples. Notably, this genetic variation is affected by various environmental factors (e.g., climate, surface type, latitude, etc.) and samples show greater diversity near the equator. Moreover, sequences associated with AMR markers are widespread, though not necessarily abundant, and show geographic specificity. Here, we present the results of our global analyses and a set of tools developed to access and analyze this extensive atlas, including: two interactive map-based visualizations for samples (metasub.org/map) and AMRs (resistanceopen.org), an indexed search tool over raw sequence data (dnaloc.ethz.ch/), a Git repository for all analytical pipelines and figures, and application programming interfaces (APIs) for computationally accessing results (github.com/metasub/metasub_utils).

## 2 Results

We have collected 4,728 samples from from the mass transit systems of 60 cities around the world (Table 1, Supplementary table S1). These samples were collected from various common surfaces in the mass transit systems such as railings, benches, and ticket kiosks and were subjected to metagenomic sequencing. We use the microbiome of mass transit systems as a proxy for the urban microbiome as a whole and present our key findings here.

**Table 1:** Sample Counts, The number of samples collected from each region.

| Region | Pilot | CSD16 | CSD17 | Other | Total |
| --- | --- | --- | --- | --- | --- |
| North America | 28 | 284 | 371 | 276 | 959 |
| East Asia | 34 | 26 | 1297 | 0 | 1357 |
| Europe | 177 | 310 | 939 | 1 | 1427 |
| Sub Saharan Africa | 0 | 116 | 192 | 0 | 308 |
| South America | 20 | 44 | 199 | 68 | 331 |
| Middle East | 0 | 100 | 15 | 0 | 115 |
| Oceania | 0 | 94 | 32 | 0 | 126 |
| Background Control | 0 | 0 | 40 | 0 | 40 |
| Lab Control | 0 | 0 | 20 | 6 | 26 |
| Positive Control | 0 | 0 | 33 | 6 | 39 |
| Total | 259 | 974 | 3138 | 357 | 4728 |

### 2.1 A Core Urban Microbiome Centers Global Diversity

We first investigated the distribution of microbial species across the global urban environment. Specifically, we asked whether the urban environment represents a singular type of microbial ecosystem or a set of related, but distinct, communities, especially in terms of biodiversity. We observed a bi-modal distribution of taxa prevalence across our dataset, which we used to define two separate sets of taxa based on the inflection points of the distribution: the putative “sub-core” set of urban microbial species that are consistently observed (>70% of samples) and the less common “peripheral” (<25% of samples) species. We also defined a set of true “core” taxa which occur in essentially all samples (>97% of samples). Applying these thresholds, we identified 1,145 microbial species (Figure 1C) that make up the sub-core urban microbiome with 31 species in the true core microbiome (Figure 1A). Core and sub-core taxa classifications were further evaluated for sequence complexity and genome coverage on a subset of samples. Of the 1,206 taxa with prevalence greater than 70%, 69 were flagged as being low quality classifications (see methods). The sub-core microbiome was principally bacterial, with just one eukaryotic taxon identified and not flagged: *Saccharomyces cerevisiae.* Notably, no archaea or viruses were identified in the group of sub-core microorganisms (note that this analysis did not include viruses newly discovered in this study). For viruses in particular, this may be affected by the sampling or DNA extraction methods used, by limitations in sequencing depth, or by missing annotations in the reference databases used for taxonomic classification, which is principally problematic with phages. It is worth noting that potentially prevalent RNA viruses are omitted with our DNA-based sampling. The three most common bacterial phyla across the world’s cities ordered by the number of species observed were *Proteobacteria, Actinobacteria,* and *Firmicutes.* To test for possible geographic bias in our data, we normalized the prevalence for each taxa by the median prevalence within each city. The two normalization methods broadly agreed (Figure ??).

**Figure 1:**
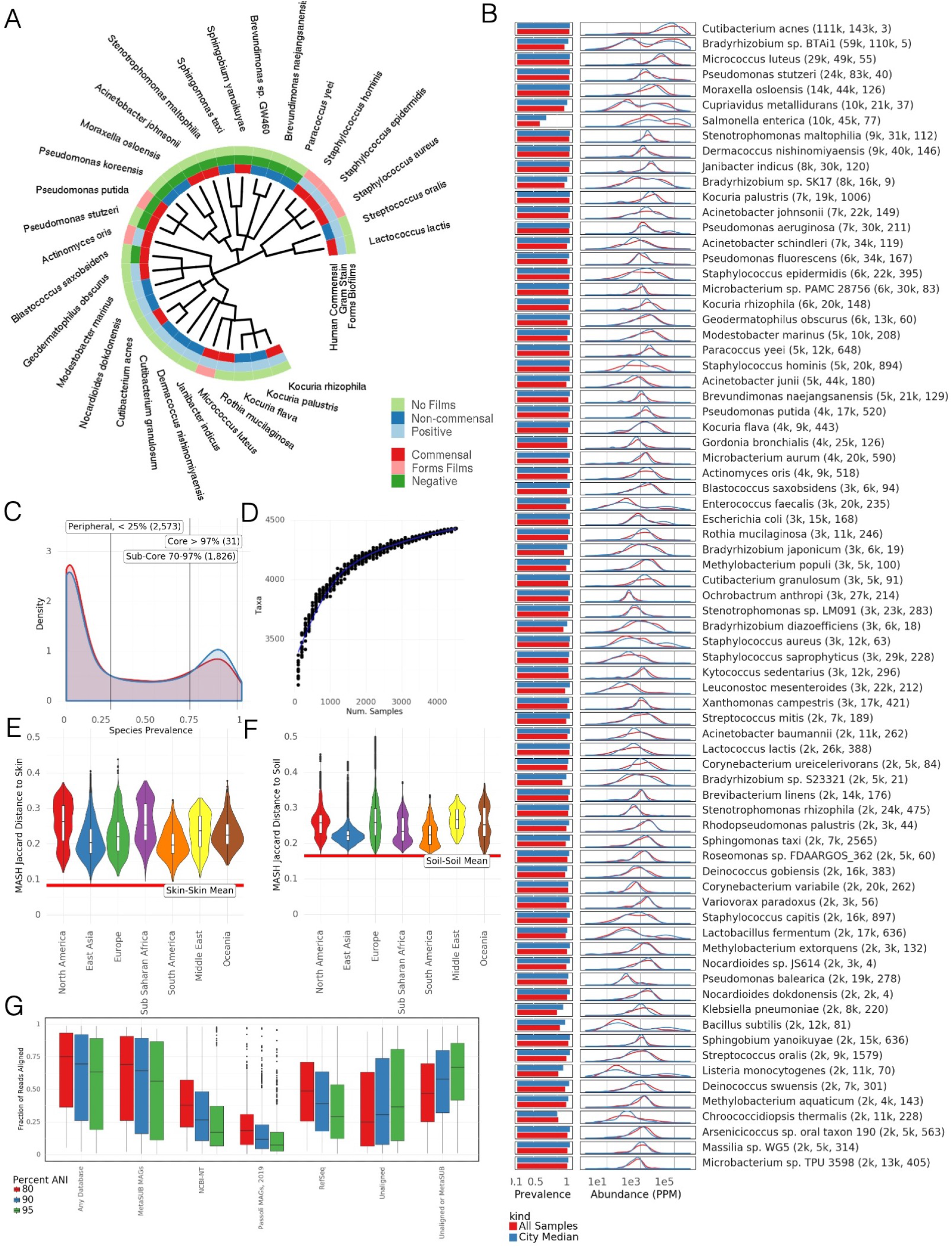
The core microbiome A) Taxonomic tree showing 31 core taxa, colored by phylum and annotated according to gram stain, ability to form biofilms, predicted association with a virus, and whether the bacteria is a human commensal species. B) prevalence and distribution of relative abundances of the 75 most abundant taxa. Mean relative abundance, standard deviation, and kurtosis of the abundance distribution are shown. C) Distribution of species prevalence from all samples and normalized by cities. Vertical lines show defined group cutoffs. D) Rarefaction analysis showing the number of species detected in randomly chosen sets of samples. E) MASH (*k*-mer based) similarity between MetaSUB samples and HMP skin microbiome samples, by continent. F) MASH (fc-mer based) similarity between MetaSUB samples and soil microbiome samples, by continent. G) Fraction of reads aligned (via BLAST) to different databases at different Average Nucleotide Identities.

Despite their global prevalence, the core taxa are not uniformly abundant across all cities. Many species exhibited a high standard deviation and kurtosis (calculated using Fisher’s definition and normal kurtosis of 0) than other species (Figure 1B). Furthermore, some species show distinctly high mean abundance, often higher than the core species, but more heterogeneous global prevalence. For example, *Salmonella enterica* is identified in less than half of all samples but is the 12th most abundant species based on the fraction of DNA that can be ascribed to it. The most relatively abundant microbial species was *Cutibacterium acnes* (Figure 1D) which had a comparatively stable distribution of abundance across all samples; *Cutibacterium acnes* is known as a prominent member of the human skin microbiome. To test for any biases arising from uneven geographic sampling, we measured the relative abundance of each taxon by calculating the fraction of reads classified to each particular taxon, and compared the raw distribution of abundance to the distribution of median abundance within each city (This process is analogous to the one used for Figure 1C, Figure 1B); the two measures closely aligned. Also, an examination of the positive and negative controls indicates that these results are not likely due to contamination or batch effect (Supp. Figure S13). In total, we observed 31 core taxa (>97%), 1,145 sub-core taxa (70-97%) 2,466 peripheral taxa (<25%), and 4,424 taxa across all samples. We term the set of all taxa observed *the urban panmicrobiome*.

To estimate the number of taxa present in our samples but which were not detected by our experimental techniques, we performed a rarefaction analysis on the taxa that were identified. By estimating the number of taxa identified for different numbers of samples, we see a diminishing trend (Figure 1D), which indicates that at some point, the species in every new sample were likely already identified in a previous one. Our rarefaction curve did not reach a plateau and, even after including all samples, it still shows an expected marginal discovery rate of roughly 1 additional species for every 10 samples added to the study. For clarity we note that this analysis only considers taxa already present in reference databases, not newly discovered taxa (below). Despite the remaining unidentified taxa, we estimate that most (80%) of the classifiable taxa in the urban microbiome could be identified with roughly 1,000 samples. However, as noted below, this new diversity is likely not evenly distributed across regions.

As humans are a major part of the urban environment, the DNA in our samples could be expected to resemble commensal human microbiomes. To investigate this, we compared non-human DNA fragments from our samples to a randomized set of 50 samples from 5 commensal microbiome sites in the Human Microbiome Project (HMP) (Consortium et al., 2012) (stool, skin, airway, gastrointestinal tract, urogenital tract). We used MASH to perform a *k*-mer based comparison of our samples vs. the selected HMP samples, which showed a roughly uniform dissimilarity between MetaSUB samples and those from different human body sites (Figure 1E, Supp. Figure S2A & B). Samples taken from surfaces that were likely to have been touched more often by human skin, such as doorknobs, buttons, railings, and touchscreens, were indeed more similar to human skin microbiomes than surfaces like bollards, windows, and the floor. Given that a large fraction of DNA in our samples could not be classified and that a *k*-mer based comparison did not find significant body-site specificity, it is possible that the unclassified DNA in our samples is from novel taxa which are not human commensals. Of note, the taxonomic composition of our samples do not closely resemble soil samples. We processed 28 metagenomic soil samples (Bahram et al., 2018) using the same pipeline as the rest of the data and compared soil samples to our samples using MASH. Our samples were very dissimilar from the soil samples (Figure 1F) even in comparison to human skin microbiomes. This suggests that the unclassified DNA may represent heretofore uncharacterized taxa that are not known commensals being shed into the environment.

We next estimated the fraction of sequences in our data that did not resemble sequences in known reference databases. We took a subset of 10,000 reads from each sample and aligned these reads to a number of reference databases using BLASTn (Altschul et al., 1990). We then identified reads that mapped to sequences in the reference databases at 80%, 90%, and 95% Average Nucleotide Identity (ANI) (Figure 1G). We used a broad set of databases for reference: RefSeq, NCBI’s NT Environmental, a large database of Metagenome Assembled Genomes (MAGs) from Pasolli et al. (2019), and MAGs from MetaSUB itself (Section 2.5). At 80% ANI, the most permissive threshold, 34.6% of reads did not map to any database while 47.3% of reads did not map or only mapped to MAGs from MetaSUB itself. This mirrors results seen by previous urban microbiome works (Afshinnekoo et al., 2015; Hsu et al., 2016).

Next, we analyzed the fraction of sequences that aligned to these same databases by region. Surprisingly, samples from Europe had the highest fraction of unaligned reads, followed by the middle east, while samples from Sub Saharan Africa had the smallest fraction of unaligned reads (Supp. Figure S1C). The proportion of reads aligned to each database did not vary significantly by region. We further investigated the relationship between geography and sample composition. In ecology, an increasing distance from the equator is associated with a decrease in taxonomic diversity (O’Hara et al., 2017). The MetaSUB data recapitulates this result and identifies a significant decrease in taxonomic diversity (though with significant noise, *p* < 2*e*16, *R*^2^ = 0.06915) as a function of absolute latitude; samples are estimated to lose 6.9672 species for each degree of latitude away from the equator (Supp. Figure S1A). The effect of latitude on species diversity is not purely monotonic, since several cities have higher species diversity then their latitude would predict. This is expected as latitude is only a rough predictor of a city’s climate. While this is an observation consistent with ecological theory, we note that our samples are heavily skewed by the location of the target cities, as well as the prevalence of those cities in specific latitude zones of the northern hemisphere.

### 2.2 Global Diversity Varies According to Covariates

Despite the core urban microbiome present in almost all samples, there was also geographic variation in taxonomy and localization. We calculated the Jaccard distance between samples measured by the presence and absence of species (which is robust to noise from relative abundance) and performed a dimensionality reduction of the data using UMAP (Uniform Manifold Approximation and Projection, McInnes et al. (2018)) for visualization (Figure 2A). Jaccard distance was correlated with distance based on Jensen-Shannon Divergence (which accounts for relative abundance) and *k*-mer distance calculated by MASH (which is based on the *k*-mer distribution in a sample, so cannot be biased by a database) (Supp. Figure S10A, B, C). In principle, Jaccard distance could be influenced by read depth as low abundance species drop below detection thresholds. However we expect this issue to be minor as the total number of species identified stabilized at 100,000 reads (Supp. Figure S9B) compared to an average of 6.01M reads per sample. Samples collected from North America and Europe were distinct from those collected in East Asia, but the separation between other regions was less clear. A similar trend was found in an analogous analysis based on functional pathways rather than taxonomy (Supp. Fig S5D), which indicates geographic stratification of the metagenomes at both the functional and taxonomic levels. Subclusters identified by UMAP roughly corresponded to city and climate but not surface type (Supp. Figure S5A, B, C). These findings confirm and extend earlier analyses performed on a fraction of the MetaSUB data which were run as a part of CAMDA Challenges in years 2017, 2018, and 2019 (camda.info).

**Figure 2:**
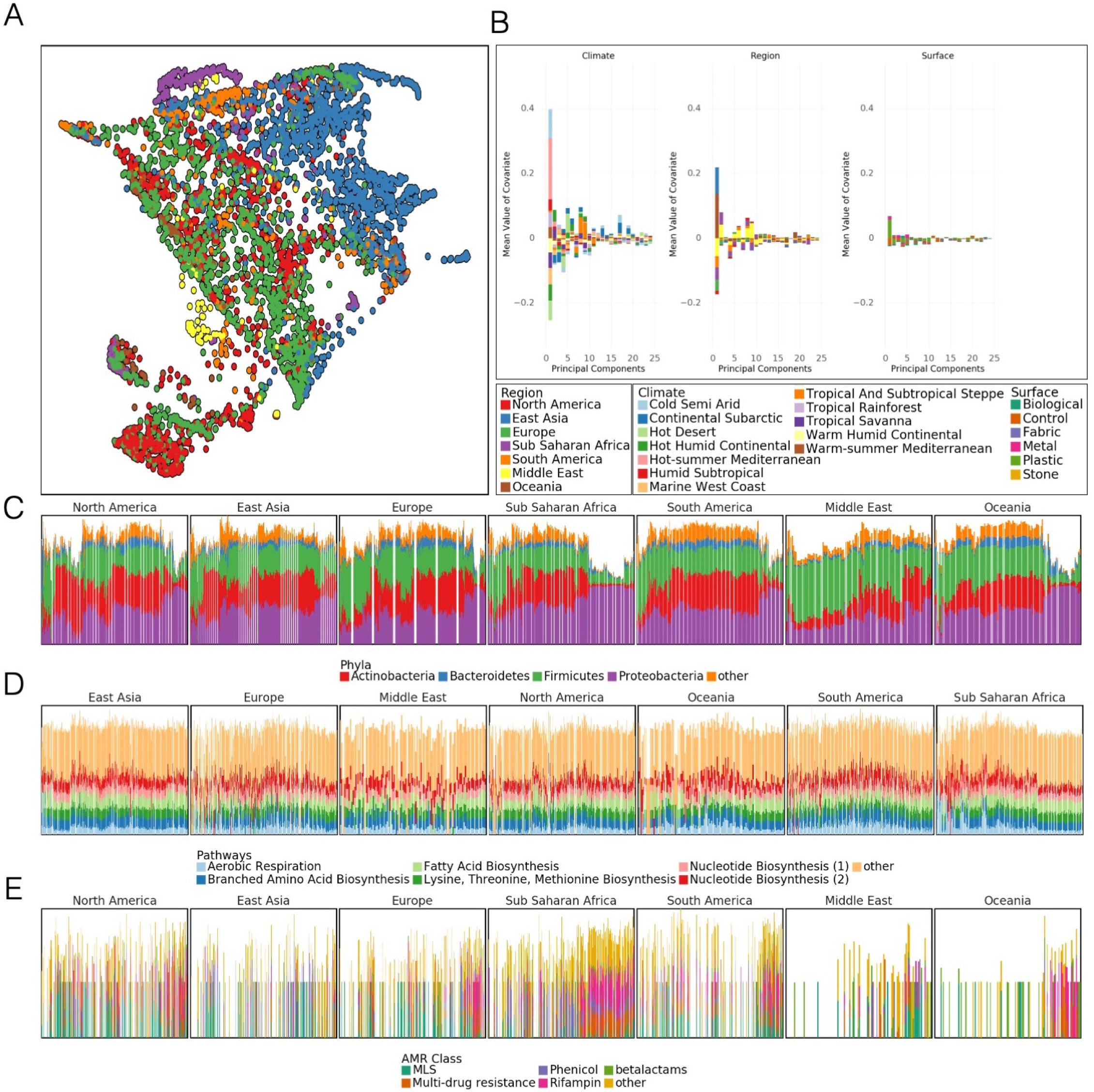
Differences at global scale A) UMAP of taxonomic profiles based on Jaccard distance between samples. Colored by the region of origin for each sample. Axes are arbitrary and without meaningful scale. The color key is shared with panel B. B) Association of the first 25 principal components of sample taxonomy with climate, continent, and surface material. C) Distribution of major phyla, sorted by hierarchical clustering of all samples and grouped by continent. D) Distribution of high-level groups of functional pathways, using the same order as taxa (C). E) Distribution of AMR genes by drug class, using the same order as taxa (C). Note that MLS is macrolide-lincosamide-streptogramin.

We quantified the degree to which metadata covariates influence the taxonomic composition of our samples using MAVRIC, a statistical tool to estimate the sources of variation in a count-based dataset (Moskowitz and Greenleaf, 2018). We identified covariates which influenced the taxonomic composition of our samples: city, population density, average temperature in June, region, elevation above sea-level, surface type, surface material, elevation above or below ground and proximity to the coast. The most important factor, which could explain 19% of the variation in isolation, was the city from which a sample was taken followed by region which explained 11%. The other four factors ranged from explaining 2% to 7% of the possible variation in taxonomy in isolation (Supp. Table S2). We note that many of the factors were confounded with one another, so they can explain less diversity than their sum. One metadata factor tested, the population density of the sampled city, had no significant effect on taxonomic variation overall.

To quantify how the principle covariates, climate, continent, and surface material impacted the taxonomic composition of samples, we performed a Principal Component Analysis (PCA) on our taxonomic data normalized by proportion and identified principal components (PCs) which were strongly associated with a metadata covariate in a positive or negative direction (PCs were centered so an average direction indicates an association). We found that the first two PCs (representing 28.0% and 15.7% of the variance of the original data, respectively) associated strongly with the city climate while continent and surface material associate less strongly (Figure 2B).

Next, we tested whether geographic proximity (in km) of samples to one another had any effect on the variation, since samples taken from nearby locations could be expected to more closely resemble one another. Indeed, for samples taken in the same city, the average JSD (Jensen-Shannon distance) was weakly predictive of the taxonomic distance between samples, with every increase of 1km in distance between two samples representing an increase of 0.056% in divergence (*p* < 2*e*16, *R*^2^ = 0.01073, Supp. Figure S1B). This suggests a “neighborhood effect” for sample similarity analogous to the effect described by Meyer et al. (2018), albeit a very minor one. To reduce bias that could be introduced by samples taken from precisely the same object we excluded all pairs of samples within 1km of one another.

At a global level, we examined the prevalence and abundance of taxa and their functional profiles between cities and continents. These data showed a fairly stable phyla distribution across samples, but the relative abundance of these taxa is unstable (Figure 2C) with some continental trends. In contrast to taxonomic variation, functional pathways were much more stable across continents, showing relatively little variation in the abundance of high level categories (Figure 2D). This pattern may also be due to the more limited range of pathway classes and their essential role in cellular function, in contrast to the much more wide-ranging taxonomic distributions examined across metagenomes. Classes of antimicrobial resistance were observed to vary by continent as well. Clusters of AMR classes were observed to occur in groups of taxonomically similar samples (Figure 2E).

We quantified the relative variation of taxonomic and functional profiles by comparing the distribution of pairwise distances in taxonomic and functional profiles. Both profiles were equivalently normalized to give the probability of encountering a particular taxon or pathway. Taxonomic profiles have a mean pairwise Jensen-Shannon Divergence (JSD) of 0.61 while pathways have a mean JSD of 0.099. The distributions of distances are significantly different (Welch’s *t*-test, unequal variances, *p* < 2*e* –16). This is consistent with observations from the Human Microbiome Project, where metabolic function varied less than taxonomic composition (Consortium et al., 2012; Lloyd-Price et al., 2017) within samples from a given body site.

### 2.3 Microbial Signatures Reveal Urban Characteristics

To facilitate characterization of novel sequences we created GeoDNA, a high-level web interface (Figure 3A) to search raw sequences against our dataset. Users can submit sequences to be processed against a *k*-mer graph-based representation of our data. Query sequences are mapped to samples and a set of likely sample hits is returned to the user. This interface will allow researchers to probe the diversity in this dataset and rapidly identify the range of various genetic sequences.

**Figure 3:**
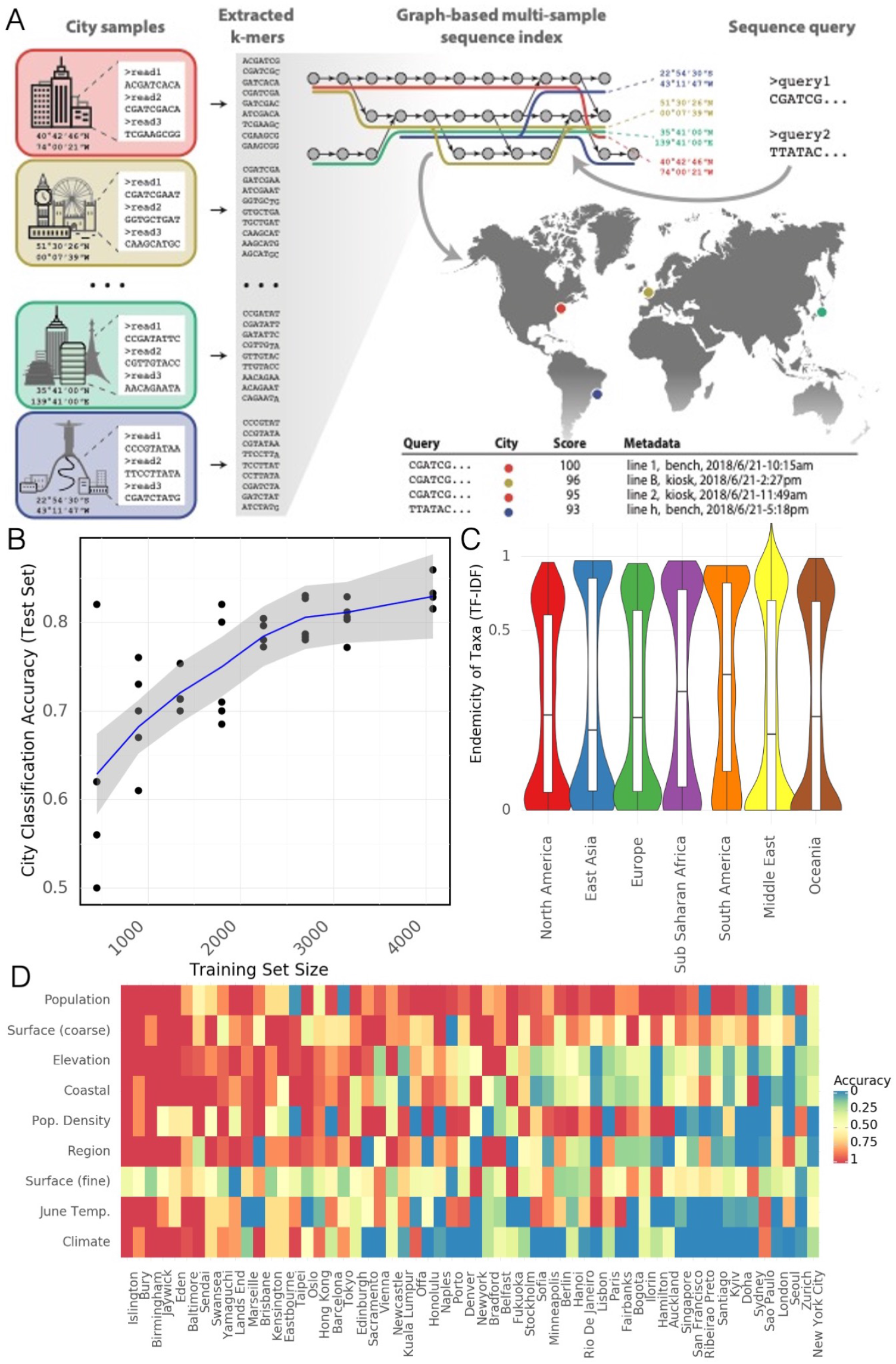
Microbial Signatures A) Schematic of GeoDNA representation generation – Raw sequences of individual samples for all cities are transformed into lists of unique fc-mers (left). After filtration, the fc-mers are assembled into a graph index database. Each *k*-mer is then associated with its respective city label and other informative metadata, such as geo-location and sampling information (top middle). Arbitrary input sequences (top right) can then be efficiently queried against the index, returning a ranked list of matching paths in the graph together with metadata and a score indicating the percentage of fc-mer identity (bottom right). The geo-information of each sample is used to highlight the locations of samples that contain sequences identical or close to the queried sequence (middle right). B) Classification accuracy of a random forest model for assigning city labels to samples as a function of the size of training set. C) Distribution of Endemicity scores (term frequency inverse document frequency) for taxa in each region. D) Prediction accuracy of a random forest model for a given feature (rows) in samples from a city (columns) that was not present in the training set. Rows and columns sorted by average accuracy. Continuous features (e.g. Population) were discretized.

We sought to determine whether a samples taxonomy reflected the environment in which it was collected. To this end we trained a Random Forest Classifier (RFC) to predict a sample’s city of origin from its taxonomic profile. We trained an RFC with 100 components on 90% of the samples in our dataset and evaluated its classification accuracy on the remaining 10%. We repeated this procedure with multiple subsamples of our data at various sizes and with 5 replicates per size to achieve a distribution (Fig. 3B). The RFC achieved 88% on held out data which compares favorably to the 7.01% that would be achieved by a randomized classifier. These results from our RFC demonstrate that city specific taxonomic signatures exist and can be predictive.

We expanded our analysis of environmental signatures in taxonomy to the prediction of features in cities not present in our training set. To do this we collated a set of 7 features for each city: population, surface material, elevation, proximity to the coast, population density, region, ave June temperature, and Koppen climate classification. We trained a RFCs to predict each feature based on all samples that were not taken from a given city then used the relevant RFC to predict the feature for samples from the held out city and recorded the classification accuracy (Figure 3D). While not all features and cities were equally predictable (in particular features for a number of British cities were roughly similar and could be predicted effectively) in general the predictions exceeded random chance by a significant margin (Supp. Figure S3A). This suggests that certain features of cities generate microbial signatures that are present globally and distinct from city specific signatures. The successful geographic classification of samples demonstrates distinct city-specific trends in the detected taxa, that may enable future forensic biogeographical capacities.

However, unique, city-specific taxa are not uniformly distributed (Figure 3B). To quantify this, we developed a score to reflect how endemic a given taxon is within a city, which reflects upon the forensic usefulness of a taxon. We define the Endemicity Score (ES) of a taxa as term-frequency inverse document frequency where the document consists of samples from some metadata defined group such as a city or region. This score is designed to simultaneously reflect the chance that a taxon could identify a given city and that that taxon could be found within the given city. A high ES for a taxon in a given city could be evidence of the evolutionary advantage that the taxon has in a particular cities environment. However, neutral evolution of microbes within a particular niche is also possible and the ES alone does not distinguish between these two hypotheses.

Note that while the ES only considers taxa which are found in a city, a forensic classifier could also take advantage of the absence of taxa for a similar metric. ES show a roughly bimodal distribution for regions (Fig. 3C). Each region possesses a number of taxa with ES scores close to 1 and a slightly larger number close to 0 (note that ES is not bounded in [0,1]). Some cities, like Offa (Nigeria), host many unique taxa while others, like Zurich (Switzerland), host fewer endemic species (Supp. Figure S3B). Large numbers of endemic species in a city may reflect geographic bias in sampling. However, some cities from well sampled continents (e.g., Lisbon, Hong Kong) also host many endemic species which would suggest that ES may indicate interchangeability and local pockets of microbiome variation for some locations.

### 2.4 Antimicrobial Resistance Genes Form Distinct Clusters

Quantification of antimicrobial diversity and AMRs are key components of global antibiotic stewardship. Yet, predicting antibiotic resistance from genetic sequences alone is challenging, and detection accuracy depends on the class of antibiotics (i.e., some AMR genes are associated to main metabolic pathways while others are uniquely used to metabolize antibiotics). As a first step towards a global survey of antibiotic resistance in urban environments, we mapped reads to known antibiotic resistance genes, using the MegaRES ontology and alignment software. We quantified their relative abundance using reads/kilobase/million mapped reads (RPKM) for 20 classes of antibiotic resistance genes detected in our samples (Figure 4A B). 2,210 samples had some sequence which were identified as belonging to an AMR gene, but no consistent core set of genes was identified. The most common classes of antibiotic resistance genes were for macrolides, lincosamides, streptogamines (MLS), and betalactams, yet the most common class of antibiotic resistance genes, MLS was found in only 56% of the samples where AMR sequence was identified.

**Figure 4:**
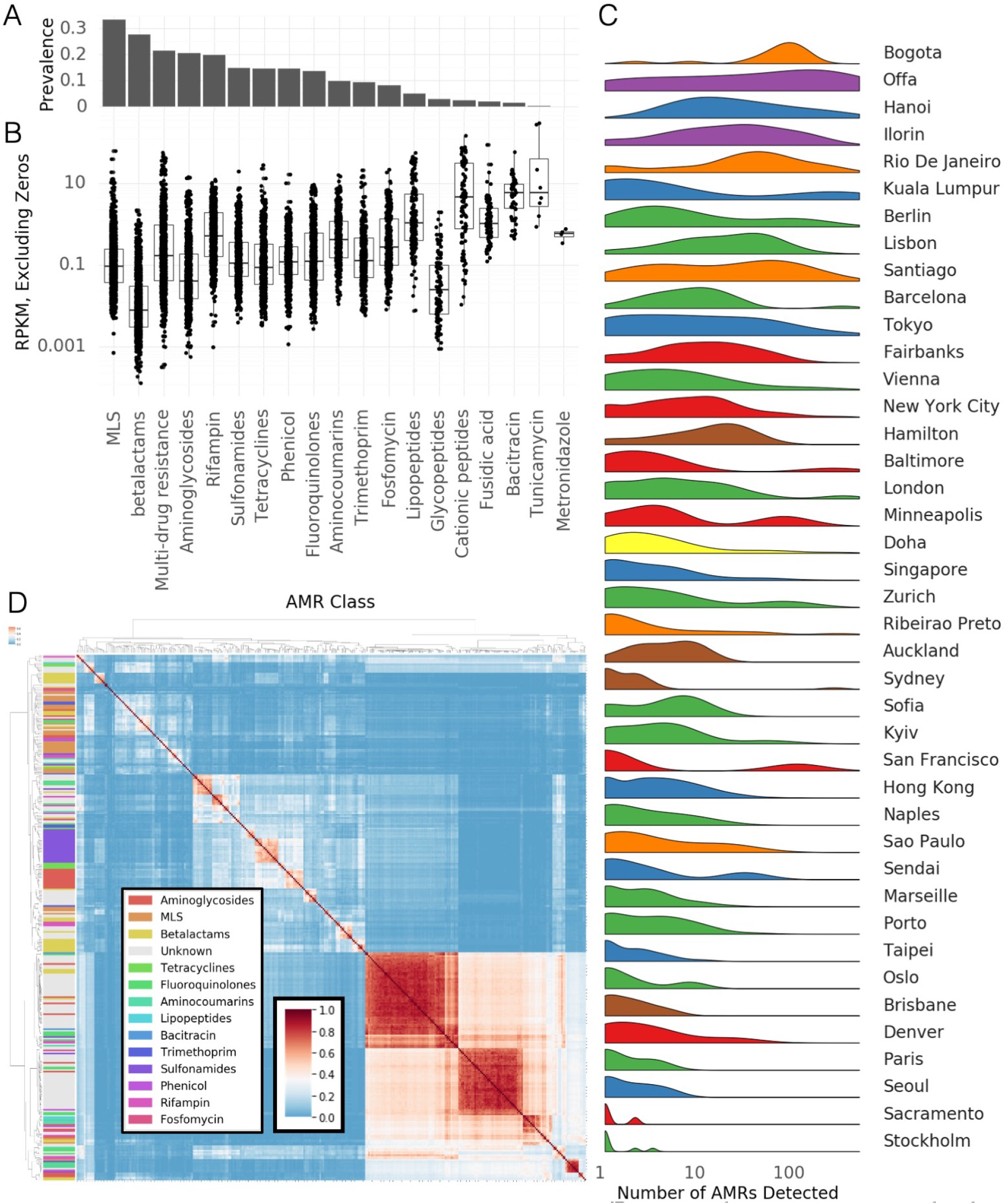
Antimicrobial Resistance Genes. A) Prevalence of AMR genes with resistance to particular drug classes. B) Abundance of AMR gene classes when detected, by drug class. C) Number of detected AMR genes by city. D) Co-occurrence of AMR genes in samples (Jaccard index) annotated by drug class.

Despite being relatively common, antibiotic resistance genes were universally in low abundance compared to functional genes, with RPKM values for resistance classes typically ranging from 0.1 – 1 compared to values of 10 – 100 for typical housekeeping genes (AMR classes contain many genes so RPKM values may be lower than they would be for individual genes). In spite of the low abundance of the genes themselves, some samples contained sequences from hundreds of distinct AMR genes. Clusters of high AMR diversity were not evenly distributed across cities (Figure 4C). Some cities had more resistance genes identified on average (15-20X) than others (e.g. Bogota) while other cities had bimodal distributions (e.g. San Francisco) where some samples had hundreds of genes while others very few. We note that 99% of the cases where we detected an AMR genes had an average depth of 2.7x, indicating that our global distribution would not dramatically change with altered read depth (Supp. Figure S6E).

As with taxa, AMR genes can be used to classify samples to cities – albeit with much less accuracy. A random forest model analogous to the one trained to predict city classification from taxonomic profiles was trained to predict from profiles of antimicrobial resistance genes. This model achieved 37.6% accuracy on held out test data (Supp. Figure S6A). While poor for actual classification this accuracy far exceeds the 7.01% that would be achieved by randomly assigning labels and indicates that there are possibly weak, city specific signatures for antimicrobial resistance genes.

Multiple AMR genes can be carried on a single plasmid and ecological competition may cause multiple taxa in the same sample to develop antimicrobial resistance. As a preliminary analysis into these phenomenons we identified clusters of AMR genes that co-occurred in the same samples (Figure 4D). We measured the Jaccard distance between all pairs of AMR genes found in at least 1% of samples and performed agglomerative clustering on the resulting distance matrix. We identified three large clusters of genes and numerous smaller clusters. Of note, these clusters often consist of genes from multiple classes of resistance. At this point we do not posit a specific ecological mechanism for this co-occurrence, but we note that the large clusters contain far more genes than are typically found on plasmids.

We performed a rarefaction analysis on the set of all resistance genes in the dataset, which we call the “panresistome” (Figure (Supp. Figure S6B). Similar to the rate of detected species, the panresistome also shows an open slope with an expected rate of discovery of 1 previously unobserved AMR gene per 10 samples. Given that AMR gene databases are rapidly expanding and that no AMR genes were found in some samples, it is likely that future analyses will identify many more resistance genes in this data.

Additionally, AMR genes show a “neighbourhood” effect within samples that are geographically proximal analogous to the effect seen for taxonomic composition (Supp. Figure S6C). Excluding samples where no AMR genes were detected, the Jaccard distance between sets of AMR genes increases with distance for pairs of samples in the same city. As with taxonomic composition. the overall effect is weak and noisy, but significant.

### 2.5 Widespread Discovery of Novel Biology

To examine these samples for novel genetic elements, we assembled and identified Metagenome Assembled Genomes (MAGs) for viruses, bacteria, and archaea and analyzed them with several algorithms. This includes thousands of novel CRISPR arrays that reflect the microbial biology of the cities and 1,304 genomes from our data, of which 748 did not match any known reference genome within 95% average nucleotide identity (ANI). 1302 of the genomes were classified as bacteria, and 2 as archaea. Bacterial genomes came predominantly from four phyla: the Proteobacteria, Actinobacteria, Firmicutes, and Bacteroidota. Novel bacteria were evenly spread across these phyla (Figure 5A).

**Figure 5:**
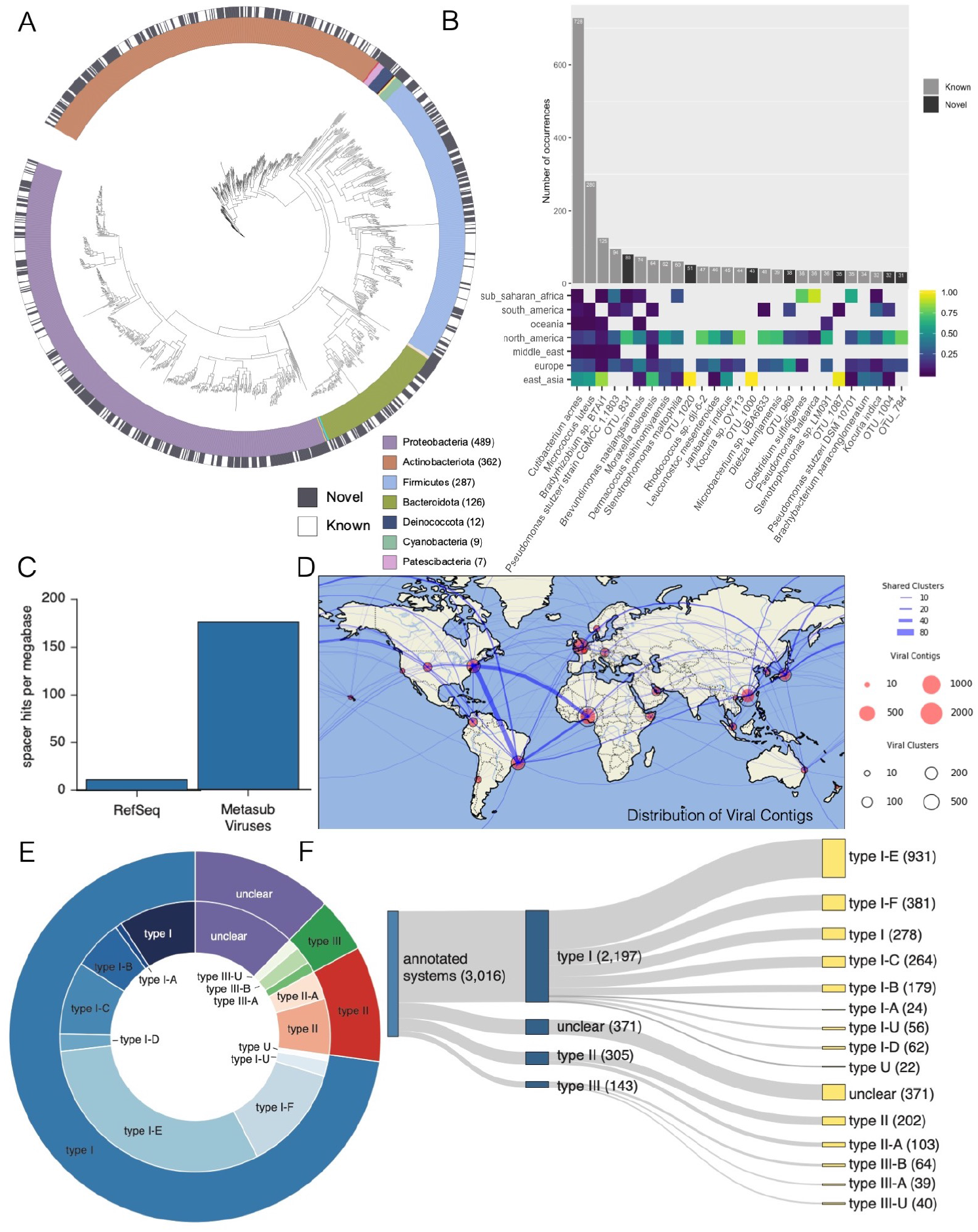
Novel Biology A) Taxonomic tree for Metagenome Assembled Genomes (MAGs) found in the MetaSUB data. Outer black and white ring indicates if the MAG matches a known species, inner ring indicates phyla of the MAG. B) Top: the number of samples where the most prevalent MAGs were found. Bottom: The regional breakdown of samples where the MAG was found. C) Mapping rate of CRISPR Spacers from MetaSUB data to viral genomes in RefSeq and viral genomes found in MetaSUB data. D) Geographic distribution of viral genomes found in MetaSUB data. E & F) Fractional breakdowns of identifiable CRISPR systems found in the MetaSUB data

Assembled bacterial genomes were often identified in multiple samples. Several of the most prevalent bacterial genomes were novel species (Figure 5B). Some assembled genomes, both novel and not, showed regional specificity while others were globally distributed. The taxonomic composition of identifiable genomes roughly matched the composition of the core urban microbiome (Section 2.1). The number of identified bacterial MAGs was somewhat based on read depth and the sample count per city (Supp. Figure S7A). The number of bacterial MAGs discovered in a city which did not match a known species was closely correlated to the total number of bacterial MAGs discovered in that city (Supp. Figure S7B). Bacterial MAGs were roughly evenly distributed geographically with the notable exception of Offa, which had dramatically more novel bacterial species than other cities.

We investigated assembled contigs from our samples to identify 16,584 predicted uncultivated viral genomes (UViGs). Taxonomic analysis of predicted UViGS to identify viral species yielded 2,009 clusters containing a total of 6,979 UViGs and 9,605 singleton UViGs for a total of 11,614 predicted viral species. Predicted viral species from samples collected within 10, 100 and 1000 kilometers of one another were agglomerated to examine their planetary distribution at different scales (Figure 5B). At any scale, most viral clusters appear to be weakly cosmopolitan; the majority of their members are found at or near one location, with a few exceptions.

We compared the predicted species to known viral sequences in the JGI IMG/VR system, which contains viral genomes from isolates, a curated set of prophages and 730k viral MAGs from other studies. Of the 11,614 species discovered in our data 94.1% did not match any viral sequence in IMG/VR (Paez-Espino et al., 2019) at the species level for a total of 10,928 novel viruses. We note that this number is surprisingly high but was obtained using a conservative pipeline (99.6% precision) and corresponded well with our identified CRISPR arrays (below). This suggests that urban microbiomes contain significant diversity not observed in other environments.

Next, we attempted to identify possible bacterial and eukaryotic hosts for our predicted viral MAGs. For the 686 species with similar sequences in IMG/VR, we projected known host information onto 2,064 MetaSUB viral MAGs. Additionally, we used CRISPR-Cas spacer matches in the IMG/M system to assign possible hosts to a further 1,915 predicted viral species. Finally, we used a database of 20 million metagenome-derived CRISPR spacers to provide further rough taxonomic assignments. Our predicted viral hosts aligned with our taxonomic profiles, 41% of species in the core microbiome (Section 2.1) had predicted viral-host interactions. Many of our viral MAGs were found in multiple locations (Figure 5D). Many viruses were found in South America, North America and Africa. Viral MAGs in Japan often corresponded to those in Europe and North America.

We identified 838,532 CRISPR arrays in our data of which 3,245 could be annotated for specific systems. The annotated CRISPR arrays were principally type 1-E and 1-F btu a number of type two and three systems were identified as well (Figure 5E, F). A number of arrays had unclear or ambiguous type assignment. Critically the spacers in our identified CRISPR arrays closely matched our predicted viral MAGs. We aligned spacers to both our viral MAGs and all viral sequences in RefSeq. The total fraction of spacers which could be mapped to our viral MAGS and RefSeq was similar (Supp. Figure S7C) but the mapping rate to our viral MAGs dramatically exceeded the mapping rate to RefSeq (Figure 5C). We present this as additional evidence supporting these novel viral MAGs.

## 3 Discussion

MetaSUB is a global network of scientists and clinicians developing knowledge of urban microbiomes by studying mass transit systems and hospitals within and between cities. We collected and sequenced 4,728 samples from 60 cities worldwide (Tables 1 and S1), constituting the first large scale metagenomic study of the urban microbiome. We also identified species that are geographically constrained and showed that these can be used to determine a samples city of origin (Section 2.2). Many of these species are associated with commensal microbiomes from human skin and airways, but we observed that urban microbiomes are nevertheless distinct from both human and soil microbiomes. Notably, no species from the *Bacteroidetes*, a prominent group of human commensal organisms (Eckburg et al., 2005; Qin et al., 2010), was identified in the core urban microbiome. We conclude that there is a consistent urban microbiome core (Figure 1, 2), which is supplemented by geographic variation (Figure 2) and microbial signatures based on the specific attributes of a city (Figure 3). Our data also indicates that significant diversity remains to be characterized and that novel taxa may be discovered in the data (Figure 5), that environmental factors affect variation, and that sequences associated with AMR are globally widespread but not necessarily abundant (Figure 4). In addition to these results, we present several ways to access and analyze our data including interactive web based visualizations, search tools over raw sequence data, and high level interfaces to computationally access results.

Unique taxonomic composition and association with covariates specific to the urban environment suggest that urban microbiomes should be treated as ecologically distinct from both surrounding soil microbiomes and human commensal microbiomes. Though these microbiomes undoubtedly interact with the urban environment, they nonetheless represent distinct ecological niches with different genetic profiles. While our metadata covariates were associated with the principal variation in our samples, they do not explain a large proportion of the observed variance. It remains to be determined whether variation is essentially a stochastic process or if a deeper analysis of our covariates proves more fruitful. We have observed that less important principal components (roughly PCs 10-100) are generally less associated with metadata covariates but that PCs 1-3 do not adequately describe the data alone. This is a pattern that was observed in the human microbiome project as well, where minor PCs (such as our Figure 2B) were required to separate samples from closely related body sites.

Much of the urban microbiome likely represents novel diversity as our samples contain a significant proportion of unclassified DNA. This finding is comparable to many other metagenomic and microbiome studies including other work done in subway environments (Afshinnekoo et al., 2015; Hsu et al., 2016), airborne microbiomes (Yooseph et al., 2013), work done by the Earth Microbiome Project (Thompson et al., 2017), and others. As noted in in Figure 1 more sensitive methodology only marginally increases the proportion of DNA that can be classified. We consider the DNA which would not be classified by a sensitive technique to be true unclassified DNA and postulate that it may derive from novel genes or species. Given that our samples did not closely resemble human commensal microbiomes or soil samples, it is possible this represents novel urban DNA sequences.

Additionally, our discovery of a large number of novel viral sequences in our data suggests that there are likely to be additional novel taxa from other domains. The fraction of predicted viral sequences which belonged to previously unobserved taxa was particularly high in our study (94.1%) however taxonomic associations of these viruses to observed microbial hosts suggests these results are not spurious. This rate of discovery may prove prescient for novel taxa in other domains, and novel discovery of taxa may help to reduce the large fraction of DNA which cannot currently be classified.

Many of the identified taxa are frequently implicated as infectious agents in a clinical setting including specific *Staphylococcus, Streptococcus, Corynebacterium, Klebsiella* and *Enterobacter* species. There is no clear indication that these species identified in the urban environment are pathogenic, and further indepth study is necessary to determine the clinical impact of urban microbiomes. This includes microbial culture studies, specifically searching for virulence factors and performing strain-level characterization. Seasonal variation also remains open to study as the majority of the samples collected here were from two global City Sampling Days (June 21, 2016 and 2017). Further studies, some generating novel data, will need to explore whether the core microbiome shifts over the course of the year, with particular interest in the role of the microbiome in flu transmission (Cáliz et al., 2018; Korownyk et al., 2018).

The COVID-19 crisis has thrown the need for broad microbial surveillance into sharp relief. Microbial genetic mapping of urban environments will give public health officials tools to assess risk, map outbreaks, and genetically characterize problematic species. This study identifies a large number of novel viruses in the environment as well as antimicrobial resistance genes in bacteria. These data will be an important starting point for mitigating future epidemics.

As metagenomics and next-generation sequencing becomes more and more available for clinical (Wilson et al., 2019) and municipal use (Hendriksen et al., 2019), it is essential to contextualize the AMR markers or presence of new species and strains within a global and longitudinal context. The most common AMR genes were found for two classes of antibiotic: MLS and beta-lactams. MLS represents macrolides, lincosamides and streptogramins, which are three groups of antibiotics with a mechanism of action of inhibiting bacterial protein synthesis. Macrolides, with strong Gram-positive and limited Gram-negative coverage, are prevalently used to treat upper respiratory, skin, soft tissue and sexually transmitted infections amongst others. Beta-lactam antibiotics are a major class of antibiotics including penicillins, cephalosporins, monobactams, carbapenems and carbacephems that are all used to treat a wide array of infections. Antimicrobial resistance has surged due to the selection pressure of widespread use of antibiotics and is now a global health issue plaguing communities and hospitals worldwide. Antimicrobial resistance genes are thought to spread from a variety of sources including hospitals, agriculture and water (Bougnom and Piddock, 2017; Klein et al., 2018). The antimicrobial classes particularly impacted by resistance include beta-lactamases, gylcopeptides and fluoroquinolones (Rice, 2012), all of which we found antimicrobial resistance genes for across our samples. We found that there was uneven distribution of AMR genes across cities. This could be the result of some of combination of different levels of antibiotic use, differences in the urban geography between cities (population density, presence of untreated wastewater etc), or reflect the background microbiome in different places in the world. Techniques to estimate antibiotic resistance from sequencing data remain an area of intense research as certain classes of AMR gene (ie. fluoroquinolones) are sensitive to small mutations and it is possible that our methods may not fully reflect true resistance. Further research is needed to fully explore AMR genes in the urban environment, including culture studies which directly measure the phenotype of resistance.

One of the challenges in the field of metagenomics of the built environment is dealing with low biomass samples. Not only does it introduce the challenge of contamination (Kim et al., 2017) which requires standardized sample preparation and the use of positive and negative controls, but there is also the challenge in biases and data interpretation (McLaren et al., 2019). Metagenomic studies rely on bioinformatics analyses that predict relative abundances of taxa, functional genes, antimicrobial resistance genes, etc. When you have low biomass samples, these relative abundances may appear high when their absolute abundance is in fact low when considering where the samples came from. However, this is an inherent component of metagenomics that studies and examines microbiomes and communities based on the metrics and measurements of relative abundances. There are important considerations to be made from sample collection to bioinformatics analysis to ensure limited biases are introduced to a study (McLaren et al., 2019). Moreover, the overall findings must be interpreted with the proper context and scope of the experiment and samples collected.

In summary, this study presents a first molecular atlas of urban and mass-transit metagenomics from across the world. By facilitating large scale epidemiological comparisons, it is a first critical step towards quantifying the clinical role of environmental microbiomes and provides requisite data for tracking changes in ecology or virulence. Moreover, in order to study the transmission of AMRs on a global scales this dataset represents only focuses on some of the sources and vectors of the built environment. Indeed, datasets from rural and suburban areas with livestock and farms, sewage from cities (Fresia et al., 2019; Joseph et al., 2019), and other notable sources of AMRs need to be integrated together to truly capture AMR mechanisms at the global scale (Singer et al., 2016; Thanner et al., 2016). Previous studies have already demonstrated a role for precision clinical metagenomics in managing infectious disease and global health (Afshinnekoo et al., 2017; Gardy and Loman, 2018; Ladner et al., 2019). As demonstrated by the coronavirus disease 2019 (COVID-19) pandemic, as an atlas this data has the potential to aid physicians, public health officers, government officials, and others in tracing, diagnosis, clinical decision making, and policy within their communities.

### 3.1 Open Science

The MetaSUB dataset is built and organized for full accessibility to other researchers. This is consistent with the concept of Open Science. Specifically, we built our study with the FAIR principles in mind: Findable, Accessible, Interoperable and Reusable.

To make our study reproducible, we released an open source version-controlled pipeline called the MetaSUB Core Analysis Pipeline (CAP). The CAP is intended to improve the reproducibility of our findings by making it easy to apply a number of analyses consistently to a large dataset. This pipeline includes all steps from extracting data from raw sequence data to producing refined results like taxonomic and functional profiles. The CAP itself is principally composed of other open peer-reviewed scientific tools, with only a few custom scripts for mundane tasks. Every tool in the CAP is open source with a permissive license. The CAP is available as a docker container for easier installation in some instances and all databases used in the CAP are available for public download. The CAP is versioned and includes all necessary databases allowing researchers to replicate results. The CAP is not designed to produce highly novel results but is meant to be a good practice agglomeration of open source tools.

However, the output of the CAP still consists of a number of different output formats with multiple files for each sample. To make our results more reproducible and accessible, we have developed a program to condense the outputs of the Core Analysis Pipeline into a condensed data-packet. This data packet contains results as a series of Tidy-style data tables with descriptions. The advantage of this set-up is that result tables for an entire dataset can be parsed with a single command in most high level analysis languages like Python and R. This package also contains Python utilities for parsing and analyzing data packets which streamlines most of the boilerplate tasks of data analysis. All development of the CAP and data packet builder (Capalyzer) package is open source and permissively licensed.

In addition to general purpose data analysis tools essentially all analysis in this paper is available as a series of Jupyter notebooks. Our hope is that these notebooks allow researchers to reproduce our results, build upon our results in different contexts, and better understand precisely how we arrived at our conclusions. By providing the exact source used to generate our analyses and figures, we also hope to be able to quickly incorporate new data or correct any mistakes that might be identified.

For less technical purposes, we also provide web-based interactive visualizations of our dataset (typically broken into city-specific groups). These visualizations are intended to provide a quick reference for major results as well as an exploratory platform for generating novel hypotheses and serendipitous discovery. The web platform used, MetaGenScope, is open source, permissively licensed, and can be run on a moderately powerful machine (though its output relies on results from the MetaSUB CAP).

Our hope is that by making our dataset open and easily accessible to other researchers the scientific community can more rapidly generate and test hypotheses. One of the core goals of the MetaSUB consortium is to build a dataset that benefits public health. As the project develops we want to make our data easy to use and access for clinicians and public health officials who may not have computational or microbiological expertise. We intend to continue to build tooling that supports these goals.

### 3.2 CAMDA

Since 2017 MetaSUB has partnered with the Critical Assessment of Massive Data Analysis (CAMDA) camda.info, a full conference track at the Intelligent Systems for Molecular Biology (ISMB) Conference. At this venue a subset of the MetaSUB data were released to the CAMDA community in the form of annual challenge addressing the issue of geographically locating samples: ‘The MetaSUB Inter-City Challenge’ in 2017 and ‘The MetaSUB Forensics Challenge’ in 2018 and 2019. In the latter challenge the MetaSUB data has been complemented by data from EMP (Thompson et al., 2017) and other studies (Delgado-Baquerizo et al., 2018; Hsu et al., 2016). This Open Science approach of CAMDA has generated multiple interesting results and concepts relating to urban microbiomics, resulting in several publications biologydirect.biomedcentral.com/articles/collections/camdaproc as well as perspective manuscript about moving towards metagenomics in the intelligence (Mason-Buck et al., 2020). The partnership is continued in 2020 with ‘The Metagenomic Geolocation Challenge’ where the MetaSUB data has been complemented by the climate/weather data in order to construct multi-source microbiome fingerprints and predict the originating ecological niche of the sample.

## 4 Data Access

Raw sequencing reads from this study contain significant amounts of human DNA and cannot yet be made public. However, reads with the majority of human DNA filtered and low quality bases removed are available for download from Wasabi (an Amazon S3 clone) with individual URLs located here: https://github.com/MetaSUB/metasub_utils. In addition to raw reads higher level results (e.g. taxonomic profiles, functional pathways, etc.) are available in the MetaSUB data packet also available for download from Wasabi. For instructional purposes we also provide a simplified data packet for teaching which includes balanced numbers of samples from each city and completely filled metadata tables.

Interactive data visualizations are available on https://pangea.gimmebio.com/contrib/metasub, https://www.metagenscope.com and GeoDNA, an interface to search query DNA sequences against MetaSUB samples, is available at (dnaloc.ethz.ch/). MetaSUB data may be downloaded from https://pangea.gimmebio.com. MetaSUB metadata is available in the data-packet, on Pangea, or may be downloaded from https://github.com/MetaSUB/MetaSUB-metadata. Programs used for analysis of data may be found at https://github.com/MetaSUB/MetaSUB_CAP and https://github.com/dcdanko/capalyzer. Jupyter notebooks used to generate the figures and statistics in this study can be found at https://www.github.com/MetaSUB/main_paper_figures. Additional tools and resources are described here https://github.com/MetaSUB/bioinformatics_management.

## 5 Acknowledgement

DCD was supported by the Tri-Institutional Training Program in Computational Biology and Medicine (CBM) funded by the NIH grant 1T32GM083937.

We thank GitHub for providing private repositories to the MetaSUB consortium at no cost.

We thank XSEDE and Philip Blood for their support of this project.

We would like to thank the Epigenomics and Genomics Core Facilities at Weill Cornell Medicine, funding from the Irma T. Hirschl and Monique Weill-Caulier Charitable Trusts, Bert L and N Kuggie Vallee Foundation, the WorldQuant Foundation, Igor Tulchinsky, The Pershing Square Sohn Cancer Research Alliance, NASA (NNX14AH50G, NNX17AB26G), the National Institutes of Health (R01ES021006, R25EB020393, 1R21AI129851, 1R01MH117406), TRISH (NNX16AO69A:0107, NNX16AO69A:0061), the NSF (1840275), the Bill and Melinda Gates Foundation (OPP1151054) and the Alfred P. Sloan Foundation (G-2015-13964), Swiss National Science Foundation grant #407540_167331 “Scalable Genome Graph Data Structures for Metagenomics and Genome Annotation” as part of Swiss National Research Programme (NRP) 75 “Big Data”

Discovery of novel viral sequences was work conducted by the US Department of Energy Joint Genome Institute, a DOE Office of Science User Facility, under contract number DE-AC02-05CH11231 and used resources of the National Energy Research Scientific Computing Center, supported by the Office of Science of the US Department of Energy.

MetaSUB Sweden was supported by Stockholm Health Authority (Region Stockholm) grant SLL 20160933 awarded to KIU.

MetaSUB Seoul was supported by the Institut Pasteur Korea (2015MetaSUB) and a National Research Foundation of Korea (NRF) grant (NRF-2014K1A4A7A01074645, 2017M3A9G6068246).

Metasub Chile was supported by funding from CONICYT Fondecyt Iniciación grant 11140666 and 11160905, as well as funding from the Millennium Science Initiative of the Ministry of Economy, Development and Tourism, Government of Chile.

MetaSUB Japan was supported by research funds from Keio University, the Yamagata prefectural government and the City of Tsuruoka.

MetaSUB Austria and Ukraine acknowledge the bilateral AT-UA collaboration fund (WTZ:UA 02/2019; Ministry of Education and Science of Ukraine, UA:M/84-2019).

MetaSUB Ukraine was supported by research funds from Kyiv Academic Univeristy, Ministry of Education and Science of Ukraine grant 0118U100290. MetaSUB Ukraine would like to express gratitude to Kyiv Metro for the support of sampling days.

MetaSUB Barcelona was supported by the Spanish Ministry of Economy and Competitiveness, ‘Centro de Excelencia Severo Ochoa 2013-2017, the CERCA Programme / Generalitat de Catalunya, the “la Caixa” Foundation, the CRG-Novartis-Africa mobility programme 2016 and TMB Director Eladio De Miguel Sainz

Work in Colombia was partially funded by Colciencias (project No. 639677758300).

Work in Sao Paulo, Brazil was partially funded by CNPq (EDN – 309973/2015-5)

Sampling was carried out in compliance with regulations and permissions from local authorities (Azienda Napoletana Mobilità s.p.a. in Naples, Italy; Régie des Transports Métropolitains in Marseille, France; Transmilenio and ANLA permit 1484 in Bogotá, Colombia; Nigerian Railway Corporation (NRC) [Ilorin and Offa Branch] and Kwara Express Transport)

We thank the many volunteers who made this study possible. Sara Abdul Majid, Natasha Abdullah, Ait-hamlat Adel, Nayra Aguilar Rojas, Affifah Saadah Ahmad Kassim, Faisal S Al-Quaddoomi, Alex Alexiev, Muhammad Al-Fath Amran, Watson Andrew, Harilanto Andrianjakarivony, Álvaro Aranguren, Carme Arnan, Freddy Asenjo, Juliette Auvinet, Nuria Aventin, Erdenetsetseg Batdelger, François Baudon, Carla Bello, Médine Benchouaia, Hannah Benisty, Anne-Sophie Benoiston, Diego Benítez, Juliana Bernardes, Tristan Bitard-Feildel, Lucie Bittner, Guillaume Blanc, Julia Boeri, Kevin Bolzli, Alexia Bordigoni, Ciro Borrelli, Sonia Bouchard, Jean-Pierre Bouly, Alessandra Breschi, Alan Briones, Aszia Burrell, Alina Butova, Dayana Calderon, Angela Cantillo, Miguel Carbajo, Katerine Carrillo, Laurie Casalot, Sofia Castro, Jasna Chalangal, Starr Chatziefthimiou, Francisco Chavez, Allaeddine Chettouh, Erika Cifuentes, Sylvie Collin, Romain Conte, Flavia Corsi, Cecilia N Cossio, Bruno D’Alessandro, Ophélie Da Silva, Katherine E Dahlhausen, Natalie R Davidson, Eleonora De Lazzari, Stéphane Delmas, Chloé Dequeker, Alexandre Desert, Clara N. Dias, Valeriia Dotsenko, Cassie L Ettinger, Emile Faure, Fazlina Fauzi, Aubin Fleiss, Juan Carlos Forero, Mathilde Garcia, Catalina García, Sonia L Ghose, Liliana Godoy, Andrea Gonzalez, Camila Gonzalez-Poblete, Charlotte Greselle, Sophie Guasco, Nika Gurianova, Sebastien Halary, Eric Helfrich, Aliaksei Holik, Chiaki Homma, Michael Huber, Stephanie Hyland, Andrea Hässig, Roland Häusler, Nathalie Hüsser, Badamnyambuu Iderzorig, Mizuki Igarashi, Shino Ishikawa, Sakura Ishizuka, Kohei Ito, Sota Ito, Tomoki Iwashiro, Marisano James, Marianne Jaubert, Marie-Laure Jerier, Guill- laume Jospin, Nao Kato, Inderjit Kaur, Akash Keluth Chavan, Mahshid Khavari, Maryna Korshevniuk, Jonas Krebs, Andrii Kuklin, Antonietta La Storia, Juliana Lago, Elodie Laine, Olha Lakhneko, Gerardo de Lamotte, Romain Lannes, Madeline Leahy, Vincent Lemaire, Dagmara Lewandowska, Manon Loubens, Olexandr Lykhenko, Salah Mahmoud, Natalka Makogon, Dimitri Manoir, German Marchandon, Natalia Marciniak, Vincent Matthys, Arif Asyraf Md Supie, Irène Mauricette Mendy, Roy Meoded, Mathilde Mignotte, Ryusei Miura, Kunihiko Miyake, Maria D Moccia, Mauricio Moldes, Jennifer Molinet, Orgil-Erdene Molomjamts, Mario Moreno, Maureen Muscat, Cristina Muñoz, Francesca Nadalin, Dorottya Nagy-Szakal, Ashanti Narce, Hiba Naveed, Thomas Neff, Wan Chiew Ng, Elsy Ngwa, Agier Nicolas, Pierre Nicolas, Abdollahi Nika, Javier Quilez Oliete, Nils Ordioni, Mitsuki Ota, Francesco Oteri, Yuya Oto, Coral Pardo-Este, Young-Ja Park, Jananan Pathmanathan, Manuel Perez, Melissa P Pizzi, María Gabriela Portilla, Leonardo Posada, Catherine E. Pugh, Kyrylo Pyrshev, Sreya Ray Chaudhuri, Hubert Rehrauer, Renee Richer, Paula Rodríguez, Paul Roldán, Sandra Roth, Maria Ruiz, Mariia Rybak, Ikuto Saito, Yoshitaka Saito, Khaliun Sanchir, Kai Sasaki, Kaisei Sato, Masaki Sato, Ryo Sato, Seisuke Sato, Yuma Sato, Oli Schacher, Christian Schori, Felipe Sepulveda, Juan C Severyn, Sarah Shalaby, Hikaru Shirahata, Jordana M Silva, Gwenola Simon, Kasia Sluzek, Rebecca Smith, Yuya Sonohara, Nicolas Sprinsky, Stefan G Stark, Chisato Suzuki, Sora Takagi, Kou Takahashi, Naoya Takahashi, Tomoki Takeda, Soma Tanaka, Andrea Tassinari, Eunice Thambiraja, Antonin Thiébaut, Takumi Togashi, Yuto Togashi, Anna Tomaselli, Itsuki Tomita, Nora C Toussaint, Takafumi Tsurumaki, Yelyzaveta Tymoshenko, Mariko Usui, Sophie Vacant, Laura E Vann, Jhovana L Velasco Flores, Fabienne Velter, Riccardo Vicedomini, Tomoro Warashina, Ayuki Watanabe, Tina Wunderlin, Olena Yemets, Tetiana Yeskova, Shusei Yoshikawa, Stas Zubenko.

## 7 Methods

### 7.1 Metadata Collection and Cleaning

Metadata from individual cities was collected from a standardized form and set of data fields. The principle fields collected were the location of sampling, the material being sampled, the type of object being sampled, the elevation above or below ground, and the station or line where the sample was collected. However, several cities were unable to use the provided apps for various reasons and submitted their metadata as separate spreadsheets. Additionally, certain metadata features, such as those related to sequencing and quality control, were added after initial sample collection.

To collate various metadata sources, we built a publicly available program which assembled a large master spreadsheet with consistent sample UUIDs. After assembling the originally collected data attributes we added normalized attributes based on the original metadata to account for surface material, control status, and features of individual cities. A full description of ontologies used is provided as part of the collating program.

### 7.2 Sample Collection and Preparation

To obtain a comprehensive picture of microbial communities within a sample it is essential to choose a sampling method which absorbs and preserves biological materials during sampling, transport and storage until DNA extraction. The effectiveness of a swab may be influenced by a number of factors, including most importantly the material of the swab tip affecting the rate at which bacteria are absorbed during the sampling process. Furthermore, the design of the transport tube and DNA preserving liquids affect the integrity of the material during transport. Finally, the amount of background contamination identified for different products should be taken into account.

#### 7.2.1 Swab Comparisons

Surface samples were collected and preserved using a flocked swab with a DNA preservation tube. Two different sets of materials were used for collection. The Copan Liquid Amies Elution Swab (ESwab, Copan Diagnostics, Cat.:480C) paired with a 1mL of Liquid Amies in a plastic, screw cap tube, referred to as ‘copan swab’ and the Isohelix Swabs (Mini-Swab, Isohelix Cat.:MS-02) referred to as ‘isohelix swabs’, which were combined with 2D Thermo Scientific™ Matrix™ storage tubes (3741-WP1D-BR/Matrix 1.0 ml/EA) referred to as ‘matrix tube’. The matrix tubes were prefilled with the preservative liquid Zymo Research DNA/RNA Shield reagent™ (R1100-250) referred to as ‘Zymo shield’. After samples were collected with Copan swabs they were transported at room temperature and stored at −80C until DNA extraction. Isohelix swabs have been stored in matrix tubes containing 400*μ*l Zymo shield preservative. Matrix tubes were also transported at room temperature and stored at −80C until DNA extraction. We tested the absorption strength of Copan and Isohelix swabs for various biological and surface materials encountered when sampling subway stations. A single surface was selected for a designated sampling area to test the absorption strength. Both swabs were moistened by submerging the swab for a few seconds in their preservative media. The area was then swabbed for 3 min. covering the selected surface. It was determined that a moistened swab would lead to greater absorption strength.

#### 7.2.2 Sampling Protocol

A standard operating procedure (SOP) was developed for the sample collection to be followed by all members of the MetaSUB consortium participating in CSD. This protocol was adapted from work by Afshinnekoo et al. (2015). The goal was to standardize as much of the sampling procedure and ensure high quality control across the various cities and sampling teams. Thus it was recommended that teams collect samples from surfaces that are present throughout most subway and transit stations and systems around the world. These included ticket kiosks, turnstiles, railings, seats or benches, etc. Some cities had to adapt the SOP according to their city especially if they did not have a subway system and were collecting samples from other transit systems. Changes to the SOP involved the types of surfaces being sampled, not the sampling procedure itself. However, the vast majority of sampling teams collected samples from these surfaces. Moreover, a significant amount of metadata was recorded throughout sample collections to ensure as much information regarding the samples was captured. All cities also developed sampling plans for their collections and submitted them for review to have swabs sent to them, this was to ensure consistency across the various sites.

All principal investigators and MetaSUB city leaders were trained in the sampling instructions and this training was further disseminated to the respective sampling teams to ensure consistent and quality control sampling. Swabbers were instructed to put on gloves before each sample collection. The swab was dipped in the preservative medium to be pre-moistened before collection and sampling was timed to 3 minutes to ensure highest yield. Sample collectors used Copan swabs in 2016 and Isohelix swabs in 2017. Other key points in training included ensure highest surface area was used for collection (i.e. swab entire bench, not just one area) and avoiding any areas that appeared wet, contaminated, and not consistent with a subway surface. Any other observations or important notes during sample collection that could add more context to data analysis and interpretation were recorded on the notes section of the metadata collection apps.

#### 7.2.3 In-Lab controls CSD2016

As positive lab control we used 30*μ*l ZymoBiOMICS Microbial Community standard (Catalog #D6300), which we added to an empty sterile urine cap, followed by swabbing with Copan Liquid Amies Elution Swab (ESwab,Copan Diagnostics, Cat.:480C) for 1.5min / 3 minutes. As negative (background) lab control we used 50*μ*l of the final resuspension buffer (MoBio PowerSoil®DNA Isolation Kit, Cat.:12888- 100), which we have added to an empty sterile urine cup followed by swabbing for 3 min (Fig. S1). Furthermore, the working space has been swabbed for 1.5 min / 3 min before and after treatment with 10% bleach (Fig. S2) to test for background contamination rates. To identify the background levels of biological material in the air at sample areas, a Copan swab has been held for 1.5 min – 3 min in the air. To estimate the source and amount of contamination in commercial swab and tube products used for MetaSUB, we tested all consumables in triplicates in the sterilized hood (UV light and 10% bleach wiped with ethanol).

#### 7.2.4 DNA Extraction and Library Preparation

Different extraction methods were benchmarked on the samples collected across 2015-2017, these included the MoBio Powersoil DNA, Promega Maxwell, and ZymoBiomics 96 MagBead kits. Samples were processed per the protocol with modifications highlighted in the Supplemental Methods. Library preparation for NGS analysis were prepped at HudsonAlpha Genome Center by the same methods as described in Afshinnekoo et al. (2015). Pilot samples collected in Barcelona and Stockholm were prepared using the QIAGEN QIAseq FX DNA Library Kit.

### 7.3 Quality Control

#### 7.3.1 Sequencing quality

We measured sequencing quality based on 5 metrics: number of reads obtained from a sample, GC content, Shannon’s entropy of *k*-mers, post PCR Qubit score, and recorded DNA concentration before PCR. The number of reads in each sample was counted both before and after quality control, we used the number of reads after quality control for our results though the difference was slight. GC content was estimated from 100,000 reads in each sample after low quality DNA and human reads had been removed. Shannon’s entropy of *k*-mers was estimated from 10,000 reads taken from each samples. PCR Qubit score and DNA concentration are described in the wet lab methods.

#### 7.3.2 Sequencing quality scores show expected trends

We measured sequencing quality based on 5 metrics: number of reads obtained from a sample, GC content (taken after removing human reads), Shannon’s entropy of *k*-mers (from 10,000 reads sampled from each sample), post PCR Qubit score, and recorded DNA concentration before PCR. We observed good separation of negative and positive controls based on both PCR Qubit and *k*-mer entropy (Supp. Figure S14). Distributions of DNA concentration and the number of reads were as expected. GC content was broadly distributed for negative controls while positive controls were tightly clustered, expected since positive controls have a consistent taxonomic profile. Comparing the number of reads before and after quality control did not reveal any major outliers.

#### 7.3.3 Batch effect appears minimal

A major concern for this low-biomass studies and large-scale studies are batch effects. The median flowcell used in our study contained samples from 3 cities and 2 continents. However, two flowcells covered 18 cities from 5 or 6 continents respectively. When samples from these flowcells were plotted using UMAP (see Section 2.2 for details) the major global trends we described were recapitulated (Supp. Figure S15A). Further, when plotting samples by PCR qubit and *k*-mer entropy (the two metrics that most reliably separated our positive and negative controls) and overlaying the flowcell used to sequence each sample only one outlier flowcell was identified and this flowcell was used to sequence a large number of background control samples (Supp. Figure S15B). Plots of the number of reads against city of origin and surface material (Supp. Figure S15C & D) showed a stable distribution of reads across cities. Analogous plots of PCR Qubit scores were less stable than the number of reads but showed a clear drop for control samples (Supp. Figure S15E & F). These results led us to conclude that batch effects are likely to be minimal.

#### 7.3.4 Strain Contamination

We used BLASTn to align nucelotide assemblies from case samples to control samples. We used a threshold of 8,000 base pairs and 99.99% identity as a minimum to consider two sequences homologous. This threshold was chosen to be sensitive without solely capturing conserved regions. We identified all connected groups of homologous sequences and found approximate taxonomic identifications by aligning contigs to NCBI-NT using BLASTn searching for 90% nucleotide identity over half the length of the longest contig in each group.

#### 7.3.5 Strain contamination is rare or absent

Despite good separation of positive and negative controls (see Section 7.3.1) we identified several species in our negative controls which were also identified as prominent taxa in the data-set as a whole (See Section 2.1). Our dilemma was that a microbial species that is common in the urban environment might also reasonably be expected to be common in the lab environment. In general, negative controls had lower *k*-mer complexity, fewer reads, and lower post PCR Qubit scores than case samples and no major flowcell specific species were observed. Similarly, positive control samples were not heavily contaminated. These results suggest samples are high quality but do not systematically exclude the possibility of contamination.

Previous studies have reported that microbial species whose relative abundance is negatively correlated with DNA concentration may be contaminants. We observed a number of species that were negatively correlated with DNA concentration (Supp. Figure S13A) but this distribution followed the same shape (but had a greater magnitude) as a null distribution of uniformly randomly generated relative abundances (Supp. Figure S13B) leading us to conclude that negative correlation may simply be a statistical artifact. We also plotted correlation with DNA concentration against each species mean relative abundance across the entire data-set (Supp. Figure S13C). Species that were negatively correlated with DNA concentration were clearly more abundant than uncorrelated species, this suggests that there may be a jackpot effect for prominent species in samples with lower concentrations of DNA but is not generally consistent with contamination.

We analyzed the total complexity of case samples in comparison to control samples. Case samples had a significantly higher taxonomic diversity (Supp. Figure S12A) than any type of negative control sample. We also compared the confidence of taxonomic assignments to control assignments for prominent taxa (Supp. Figure S12B) using the number of unique marker *k*-mers to compare assignments. We found that case samples had more and higher quality assignments than could be found in controls. One species, *Bradyrhizobium sp. BTAi1,* was not clearly better in case samples than controls but in this case we were able to assemble genomes for this species in several unique samples so we feel it is ambiguous.

Finally, we compared assemblies from negative controls to assemblies from our case samples searching for regions of high similarity that could be from the same microbial strain. We reasoned that uncontaminated samples may contain the same species as negative controls but were less likely to contain identical strains. Only 137 case samples were observed to have any sequence with high similarity to an assembled sequence from a negative control (8,000 base pairs minimum of 99.99% identity). The identified sequences were principally from *Bradyrhizobium* and *Cutibacterium*. Since these genera are core taxa (See Section 2.1) observed in nearly every sample but high similarity was only identified in a few samples, we elected not to remove species from these genera from case samples.

#### 7.3.6 K-Mer Based Analyses

We generated 31-mer profiles for raw reads using Jellyfish. All *k*-mers that occurred at least twice in a given sample were retained. We also generated MASH sketches from the non-human reads of each sample with 10 million unique minimizers per sketch.

We calculated the Shannon’s entropy of *k*-mers by sampling 31-mers from a uniform 10,000 reads per sample. Shannon’s entropy of taxonomic profiles was calculated using the CAPalyzer package (Section 4).

#### 7.3.7 K-Mer based metrics correlate with taxonomic metrics

We found clear correlations between three pairwise distance metrics (Supp. Figure S10A, B, C): *k*-mer based Jaccard distance (MASH), taxonomic Jaccard distance, and taxonomic Jensen-Shannon divergence. This suggests that taxonomic variation reflects meaningful variation in the underlying sequence in a sample.

We also compared alpha diversity metrics (Supp. Figure S10D): Shannon entropy of *k*-mers, and Shannon entropy of taxonomic profiles. As with pairwise distances these metrics were correlated though noise was present. This noise may reflect sub-species taxonomic variation in our samples.

#### 7.3.8 Sequence Preprocessing

Sequence data were processed with AdapterRemoval (v2.17, Schubert et al. (2016)) to remove low quality reads and reads with ambiguous bases. Subsequently reads were aligned to the human genome (hg38, including alternate contigs) using Bowtie2 (v2.3.0, fast preset, Langmead and Steven L Salzberg (2013)). Read pairs where both ends mapped to the human genome were separated from read pairs where neither mate mapped. Read pairs where only one mate mapped were discarded. Hereafter, we refer to the read sets as human reads and non-human reads.

#### 7.3.9 Unmapped DNA is not similar to any known sequence

A large proportion of the reads in our samples were not mapped to any references sequences. There are three major reasons why a fragment of DNA would not be classified in our analysis 1) The DNA originated from a non-human and non-microbial species which would not be present in the databases we used for classification 2) Our classifier (KrakenUniq) failed to classify a DNA fragment that was in the database due to slight mismatch 3) The DNA fragment is novel and not represented in any existing database. Explanations (1) and (2) are essentially drawbacks of the database and computational model used, and we can quantify them by mapping reads using a more sensitive aligner to a larger database, such as BLASTn (Altschul et al., 1990), or ensemble methods for analysis (McIntyre et al., 2017). To estimate the proportion of reads which could be assigned, we took 10k read subsets from each sample and mapped these to a set of large database using BLASTn (see 2.1 for details). This resulted in 34.6% reads which could not be mapped to any external database compared to 41.3% of reads mapped using our approach with KrakenUniq. We note that our approach to estimate the fraction of reads that could be classified using BLASTn does not account for hits to low quality taxa which would ultimately be discarded in our pipeline, and so represents a worst-case comparison. Explanation (3) is altogether more interesting and we refer to this DNA as true unclassified DNA. In this analysis we do not seek to quantify the origins of true unclassified DNA except to postulate that it may derive from novel species as have been identified in other similar studies.

### 7.4 Computational Analysis

We processed raw reads from all samples into taxonomic, functional and AMR profiles for each sample using the MetaSUB Core Analysis Pipeline (Danko and Mason, 2020) (v1.0.0). This includes a preprocessing stage that consists of AdapterRemoval (Schubert et al., 2016) and Human sequence removal with Bowtie2 (Langmead and Steven L Salzberg, 2013). Pre-processed reads were subsequently processed according to the methods below.

#### 7.4.1 Taxonomic Analysis

We generated taxonomic profiles by processing non-human reads with KrakenUniq (v0.3.2 Breitwieser et al. (2018)) using a database based on all draft and reference genomes in NCBI/RefSeq Microbial (bacteria/archaea, fungi and virus) ca. March 2017. KrakenUniq was selected because its high performance, as it has been demonstrated to be comparable or having higher sensitivity than the best tools identified in a recent benchmarking study (McIntyre et al. (2017)) on the same comparative dataset. In addition, KrakenUniq allows for tunable specificity and identifies *k*-mers that are unique to particular taxa in a database. Reads are broken into *k*-mers and searched against this database. Finally, the taxonomic makeup of a sample is given by identifying the taxa with the greatest leaf to ancestor weight.

KrakenUniq reports the number of unique marker *k*-mers assigned to each taxon, as well as the total number of reads, the fraction of available marker *k*-mers found, and the mean copy number of those *k*-mers. We found that requiring more *k*-mers to identify a species resulted in a roughly linear decrease in the total number of species identified without a plateau or any other clear point to set a threshold (Supp. Figure S9A). In an ongoing but unpublished clinical study we have used a threshold of 512 marker *k*-mers to accurately recapitulate the results of culturing while identifying few species which were not cultured. Since false positives are less problematic in the current study than in a clinical study and because we could use our large number of samples as a partially orthogonal confirmation we chose less strict thresholds for KrakenUniq in this study.

At a minimum we required three reads assigned to a taxa with 64 unique marker *k*-mers. This setting captures a group of taxa with low abundance but reasonable (~ 10-20%) coverage of the *k*-mers in their marker set (Supp. Figure S9C). However, this also allows for a number of taxa with very high (105) duplication of the identified marker *k*-mers and very few *k*-mers per read which we believe is biologically implausible (Supp. Figure S9D). We filtered these taxa by applying a further filter which required that the number of reads not exceed 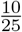 times the number of unique *k*-mers, unless the set of unique *k*-mers was saturated (> 90% completeness). We include a full list of all taxonomic calls from all samples including diagnostic values for each call. We do not attempt to classify reads below the species level in this study.

We further evaluated prominent taxonomic classifications for sequence complexity and genome coverage. For each microbe evaluated we calculated two indices generated using a random subset of 152 samples: the average topological entropy of reads assigned to the microbe and the Gini-coefficient of read positions on the microbial genome. For brevity we refer to these as *mean sequence entropy* (MSE) and *coverage equality* (CE). The formula for topological entropy of a DNA sequence is described by Koslicki (2011). Values close to 0 correspond to low-complexity sequences and values near 1 are high complexity. In this work we use a word size of 3 with an overall sequence length of 64 since this readily fits into our reads. To find the MSE of a microbial classification we take the arithmetic mean of the topological entropy of all reads that map to a given microbial genome in a sample. The Gini-coefficient is a classic economic measure of income inequality. We repurpose it here to evaluate the evenness of read coverage over a microbial classification. Reads mapping to a microbial genome are assigned to a contiguous 10kbp bin and the Gini-coefficient of all bins is calculated. Like MSE, the Gini-coefficient is bounded in [0,1]. Lower values indicate greater inequality, very low values indicate that a taxon may be misidentified from conserved and near conserved regions. We downloaded one representative genome per species evaluated and mapped all reads from samples to using Bowtie2 (sensitive-local preset). Indices were processed from alignments using a custom script. Species classifications with an average MSE less than 0.75 or CE less than 0.1 were flagged.

To determine relative abundance of taxa where applicable we rarefied samples to 100,000 classified reads, computed the proportion of reads assigned to each taxon, and took the distribution of values from all samples. This was the minimum number of reads sufficient to maintain taxonomic richness (Supp. Figure S9B). We chose sub-sampling (sometimes referred to as rarefaction in the literature) based on the study by Weiss et al. (2017), showing that sub-sampling effectively estimates relative abundance. Note that we use the term prevalence to describe the fraction of samples where a given taxon is found at any abundance and we use the term relative abundance to describe the fraction of DNA in a sample from a given taxon.

We compared our samples to metagenomic samples from the Human Microbiome Project and a metagenomic study of European soil samples using MASH (Ondov et al., 2016), a fast *k*-mer based comparison tool. We built MASH sketches from all samples with 10 million unique *k*-mers to ensure a sensitive and accurate comparison. We used MASH’s built-in Jaccard distance function to generate distances between our samples and HMP samples. We then took the distribution of distances to each particular human commensal community as a proxy for the similarity of our samples to a given human body site.

We also compared our samples to HMP and soil samples using taxonomic profiles generated by MetaPhlAn v2.0 (Segata et al., 2012). We generated taxonomic profiles from non-human reads using MetaPhlAn v2.0 and found the cosine similarity between all pairs of samples.

We used the Microbe Directory (Shaaban et al., 2018) to annotate taxonomic calls. The Microbe Directory is a hand curated, machine readable, database of functional annotations for 5,000 microbial species.

#### 7.4.2 Functional Analysis

We analyzed the metabolic functions in each of our samples by processing non-human reads with HUMAnN2 (Franzosa et al., 2018). We aligned all reads to UniRef90 using DIAMOND (v0.8.36, (Buchfink et al., 2014)) and used HUMAnN2 to produce estimate of pathway abundance and completeness. We filtered all pathways that were less than 50% covered in a given sample but otherwise took the reported pathway abundance as is after relative abundance normalization (using HUMAnN2’s attached script).

High level categories of functional pathways were found by grouping positively correlated pathways and manually annotating resulting clusters.

### 7.5 Assembly and Plasmid Annotations

All samples were assembled using metaSPAdes (v3.8.1 Nurk et al. (2017)) with default settings. Assembled scaffolds of at least 1,500bp of length were annotated using PlasFlow (v1.1 Krawczyk et al. (2018)) using default settings. PlasFlow predicts whether a contig is likely from a chromosome or a plasmid and gives a rough taxonomic annotation. Predicting which sequences are from plasmids is a difficult problem and some annotations may be incorrect.

#### 7.5.1 Analysis of Antimicrobial Resistance Genes

We generated profiles of antimicrobial resistance genes using MegaRes (v1.0.1, Lakin et al. (2017)). To generate profiles from MegaRes, we mapped non-human reads to the MegaRes database using Bowtie2 (v2.3.0, very-sensitive presets, Langmead and Steven L Salzberg (2013)). Subsequently, alignments were analyzed using ResistomeAnalyzer (commit 15a52dd github.com/cdeanj/resistomeanalyzer) and normalized by total reads per sample and gene length to give RPKMs. MegaRes includes an ontology grouping resistance genes into gene classes, AMR mechanisms, and gene groups. AMR detection remains a difficult problem and we note that detection of a homologous sequence to a known AMR gene does not necessarily imply an equivalent resistance in our samples. Currently, the gold standard for detecting AMR is via culturing.

Known AMR genes can come from gene families with homologous regions of sequence. To reduce spurious mapping from gene homology we used BLASTn to align all MegaRes AMR genes against themselves. We considered any connected group of genes with an average nucleotide identity of 80% across 50% of the gene length as a set of potentially confounded genes. We collapsed all such groups into a single pseudo-gene with the mean abundance of all constituent genes. Before clustering genes we removed all genes which were annotated as requiring SNP verification to predict resistance.

In addition to MegaRes we mapped non-human reads from all samples to the amino acid gene sequences in the Comprehensive Antibiotic Resistance Database (McArthur et al., 2013) using DIAMOND. While we do not use this analysis explicitly in this study we provide the results as a data table.

Assembled contigs were annotated for AMR genes using metaProdigal (Hyatt et al., 2010), HMMER3 (Eddy, 2011), and ResFam (Gibson et al., 2015) as described by Rahman et al. (2018). All predicted gene annotations with an e-value higher than 10^-10^ were discarded.

#### 7.5.2 Beta Diversity

Inter-sample (beta) diversity was measured by using Jaccard distances. We note that Jaccard distances do not use relative abundance information. Matrices of Jaccard distances were produced using built in SciPy functions treating all elements greater than 0 as present. Hierarchical clustering (average linkage) was performed on the matrix of Jaccard distances using SciPy (https://www.scipy.org/).

Dimensionality reduction of taxonomic and functional profiles was performed using UMAP (McInnes et al., 2018) on the matrix of Jaccard distances with 100 neighbours (UMAP-learn package, random seed of 42). We did not use Principal Component Analysis as a preprocessing step before UMAP as is sometimes done for high dimensional data.

#### 7.5.3 Alpha Diversity

Intra-sample (alpha) diversity was measured by using Species Richness and Shannon’s Entropy. We took species richness as the total number of detected species in a sample after rarefaction to 1 million reads. Shannon’s entropy is robust to sample read depth and accounts for the relative size of each group in diversity estimation. Shannon’s entropy is typically defined as *H* = Σ*a_i_log_2_a_i_* where *a_i_* is the relative abundance of taxon *i* in the sample. For alpha diversity based on *k*-mers or pathways, we simply substitute the relative abundance of a species for the relative abundance of the relevant type of object.

#### 7.5.4 GeoDNA Sequence Search

For building the sequence graph index, each sample was processed with KMC (version 3, [1]) to convert the reads in FASTA format into lists of *k*-mer counts, using different values of k ranging from 13 to 19 in increments of 2. All *k*-mers that contained the character “N” or occurred in a sample less than twice were removed. For each value of k, we built a separate index, consisting of a labeled de Bruijn graph, using an implicit representation of the complete graph and a compressed label representation based on Multiary Binary Relation Wavelet Trees (Multi-BRWT). For further details, we refer to the manuscript [2]. To build the index, for each sample the KMC *k*-mer count lists were transformed into de Bruijn graphs, from which path covers in the form of contig sets were extracted and stored as intermediate FASTA files. The contig sets of each sample were then transformed into annotation columns (one column per sample) by mapping them onto an implicit complete de Bruijn graph of order k. All annotation columns were then merged into a joint annotation matrix and transformed into Multi-BRWT format. Finally, the topology of the Multi-BRWT representation was optimized by relaxing its internal tree arity constraints to allow for a maximum arity of 40.

### 7.6 Novel Biology

### 7.7 Identifying Bacteria and Archaea

#### Metagenomic Assembly and Binning

All samples were re-assembled with metaSPAdes (v3.10.1 Nurk et al., 2017); generated contigs with length <1000nt were excluded from further analysis. Remaining contigs were binned with MetaBAT2 (v2.12.1 Kang et al. (2019)) with default parameters, resulting in 14,080 bins. As MetaBAT2 uses contig abundance (mean base coverage) in its analysis, we mapped reads back to their respective contigs via Bowtie2 (v2.3.4.1 Langmead and Steven L Salzberg (2013))with the flags -local -very-sensitive-local to provide accurate coverage metrics. Draft genome quality was assessed via CheckM (v1.0.13 Parks et al. (2015)) lineage_wf workflow with default parameters. Using the strategy proposed by Parks et al. (2018) we filtered bins by quality score, defined as QS = completeness – 5 * contamination; bins with QS < 50 were removed from consideration. The remaining 6,107 bins were labeled by quality based on the MIMAG standard (Bowers et al. (2018)), with some modification: 1,448 high quality (completeness >90%, contamination <5%, strain heterogeneity <0.5%) bins, 4,532 medium quality (completeness >50%, contamination <5%) bins, all others low quality. Bins of at least medium quality were selected as acceptable MAGs (5,980 total).

#### MAG Dereplication

OTUs (MAG representatives) were chosen with a two-step clustering strategy. Single-linkage clustering formed primary clusters of MAGs based on Mash ANI (v2.1.1), with intra-cluster identity at 90%. Though Mash ANI can be inaccurate for potentially incomplete genomes (Olm et al. (2017)), we can leverage the technique’s speed for the many pairwise comparisons needed in this granular step. Within primary clusters, MAGs were compared pairwise by a more accurate whole-genome ANI (gANI) via dnadiff (v1.3) from MUMmer (v3.23 Kurtz et al. (2004)). Secondary, more refined clusters were grouped based n gANI using average-linkage hierarchical clustering from the R package dendextend (v1.12.0 Galili (2015)). A gANI cut-off of 95% resulted in 1,304 representative OTUs.

#### OTU to Reference Genome Matching

OTUs were compared against reference genomes from Ref- Seq (release 96 from November 2019, complete bacterial and archaeal genomes only, with “Exclude anomalous” and “Exclude derived from surveillance project” applied) as well as the full Integrated Gut Genomes (IGG) dataset (v1.0 Nayfach et al. (2019); 23,790 representative genomes). A MinHash sketch was created for each reference genome via Mash (v2.1.1) with default parameters to find Mash distances and select candidate “best matches” from each reference database. Then, dnadiff (v1.3) was used to further quantify differences between each OTU and its best match from either database. ANI between OTUs and their matches was found as “M-to-M AvgIdentity” in the query report column (ANI 95% over 60% OTU sequence qualified as a match).

#### OTU Taxonomic Assignment

OTUs were placed into a bacterial or archaeal reference tree (based on the Genome Database Taxonomy, GTDB) and then assigned taxonomic classifications using GTDB-Tk (v1.0.2 Chaumeil et al. (2019)). GTDB-Tk relies on 120 bacterial and 122 archaeal marker genes; domain assignment is chosen based on domain-specific marker content of the OTU sequence. Using the GTDB-Tk placements, we built an OTU-only bacterial phylogeny with FastTree (v2.1.10 Price et al. (2010)). The tree was visualized using iTOL (v5.5 Letunic and Bork (2019)).

#### 7.7.1 Viral Discovery

We followed the protocol described by Paez-Espino et al. (2017). Briefly, we used an expanded and curated set of viral protein families (VPFs) as bait in combination with recommended filtering steps to identify 16,584 UViGs directly from all MetaSUB metagenomic assemblies greater than 5kb. Then, the UViGs were clustered with the content of the IMG/VR system (a total of over 730k viral sequences including isolate viruses, prophages, and UViGs from all kind of habitats). The clustering step relied on a sequence-based classification framework (based on 95% sequence identity across 85% of the shortest sequence length) followed by the markov clustering (mcl). This approach yielded 2,009 viral clusters (ranging from 2-611 members) and 9,605 singletons (or viral clusters of 1 member), sequences that failed to cluster with any sequence from the dataset or the references from IMG/VR, resulting in a total of 11,614 vOTUs. We define viral species from vOTUs as sequences sharing at least 95% identity over 85% of their length. Out of this total MetaSUB viral diversity, only 686 vOTUs clustered with any known viral sequence in IMG/VR.

#### 7.7.2 Identifying Host Virus Interactions

We used two computational methods to reveal putative host-virus connections (Paez-Espino et al., 2016a). (1) For the 686 vOTUs that clustered with viral sequences from the IMG/VR system, we projected the known host information to all the members of the group (total of 2,064 MetaSUB UViGs). (2) We used bacterial/archaeal CRISPR-Cas spacer matches (from the IMG/M 1.1 million isolate spacer database) to the UViGs (allowing only for 1 SNP over the whole spacer length) to assigned a host to 1,915 MetaSUB vOTUs. Additionally, we also used a database of over 20 million CRISPR-Cas spacers identified from metagenomic contigs from the IMG/M system with taxonomy assigned. Since some of these spacers may derive from short contigs these results should be interpreted with caution.

#### 7.7.3 CRISPR Array Detection and Annotation

Using CRISPRCasFinder the MetaSUB database was investigated to predict CRISPR arrays and annotate them with their corresponding predicted type based on CRISPR-Cas genes in their vicinity. CRISPRCasFinder was run with default parameters, “-so” and “-cas” options to identify cas genes. The precision and recall of the virus detection was 99.6% and 37.5% respectively, as previously reported by (Paez-Espino et al., 2016b).

CRISPR-Cas types were assigned to arrays based on detected cas genes within a 10 kilobases vicinity. Cases where CRISPRCasFinder associated several cas genes of contradicting CRISPR-Cas types with the same CRISPR array were regarded as unclear annotation. This procedure yielded 838,532 predicted CRISPR arrays (with additional CRISPR arrays predicted with default parameters for PILER-CR), of which, 3,245 CRISPR arrays had unambiguous annotation, resulting in 43,656 unique spacers queried against genomic databases using BLASTN.

### 7.8 Organisms/BLAST Databases

In order to associate detected spacers within defined groups (plasmids, prophages, viruses) four different genomic databases were aggregated to be searched with BLASTN. The aggregated database consisted of IMG/VR, PHASTER, and PLSDB alongside bacterial and archaeal genomic sequences from the National Center for Biotechnology Information (NCBI). All database downloads were made on the 28th January 2020. Detected and annotated spacers were searched against the databases mentioned above using BLASTN with the following additional arguments, which correspond to the default parameters of CRISPRTarget: word_size=7, evalue=1, gapopen=10, gapextend=2, penalty=-1, reward=1.

### 7.9 MetaSUB Genomic Database and Statistical Analysis

Genomic data was acquired from the MetaSUB database and matched by sample names to the corresponding metadata downloaded from the MetaSUB-metadata github repository (https://github.com/MetaSUB/MetaSUB-metadata). All data derived from MetaSUB and the subsequent steps described above was then analysed using Python 3.6. Python packages plotnine, plotly, matplotlib and seaborn where used for plotting as well as pandas to create and manage dataframes. The heatmap is clustered by Euclidean distance on the columns.

## 8 Contributing Members of the MetaSUB Consortium

Marcos Abraao, Muhammad Afaq, Ireen Alam, Gabriela E Albuquerque, Kalyn Ali, Lucia E Alvarado-Arnez, Sarh Aly, Jennifer Amachee, Maria G. Amorim, Majelia Ampadu, Nala An, Núria Andreu Somavilla, Michael Angelov, Verónica Antelo, Catharine Aquino, Mayra Arauco Livia, Luiza F Araujo, Jenny Arevalo, Lucia Elena Alvarado Arnez, Fernanda Arredondo, Matthew Arthur, Sadaf Ayaz, Silva Baburyan, Abd-Manaaf Bakere, Katrin Bakhl, Thais F. Bartelli, Kevin Becher, Joseph Benson, Denis Bertrand, Silvia Beurmann, Christina Black, Brittany Blyther, Bazartseren Boldgiv, Gabriela P Branco, Christian Brion, Paulina Buczansla, Catherine M Burke, Irvind Buttar, Jalia Bynoe, Sven Bönigk, Kari O Bøifot, Hiram Caballero, Alessandra Carbone, Anais Cardenas, Ana V Castro, Ana Valeria B Castro, Astred Castro, Simone Cawthorne, Jonathan Cedillo, Salama Chaker, Allison Chan, Anastasia I Chasapi, Gregory Chem, Jenn-Wei Chen, Michelle Chen, Xiaoqing Chen, Ariel Chernomoretz, Daisy Cheung, Diana Chicas, Hira Choudhry, Carl Chrispin, Kianna Ciaramella, Jake Cohen, David A Coil, Colleen Conger, Ana F. Costa, Delisia Cuebas, Aaron E Darling, Pujita Das, Lucinda B Davenport, Laurent David, Gargi Dayama, Paola F De Sessions, Chris K Deng, Monika Devi, Felipe S Dezem, Sonia Dorado, LaShonda Dorsey, Steven Du, Alexandra Dutan, Naya Eady, Stephen Eduard Boja Ruiz, Jonathan A Eisen, Miar Elaskandrany, Lennard Epping, Juan P Escalera-Antezana, Iqra Faiz, Luice Fan, Nadine Farhat, Kelly French, Skye Felice, Laís Pereira Ferreira, Gabriel Figueroa, Denisse Flores, Marcos AS Fonseca, Jonathan Foox, Aaishah Francis, Pablo Fresia, Jacob Friedman, Jaime J Fuentes, Josephine Galipon, Laura Garcia, Annie Geiger, Samuel M Gerner, Dao Phuong Giang, Matías Giménez, Donato Giovannelli, Dedan Githae, Samantha Goldman, Gaston H Gonnet, Juana Gonzalez, Irene González Navarrete, Tranette Gregory, Felix Hartkopf, Arya Hawkins-Zafarnia, Nur Hazlin Hazrin-Chong, Tamera Henry, Samuel Hernandez, David Hess-Homeier, Yui Him Lo, Lauren E Hittle, Nghiem Xuan Hoan, Irene Hoxie, Elizabeth Humphries, Shaikh B Iqbal, Riham Islam, Sharah Islam, Takayuki Ito, Tomislav Ivankovic, Sarah Jackson, JoAnn Jacobs, Esmeralda Jiminez, Ayantu Jinfessa, Takema Kajita, Amrit Kaur, Fernanda de Souza Gomes Kehdy, Vedbar S Khadka, Shaira Khan, Michelle Ki, Gina Kim, Hyung Jun Kim, Sangwan Kim, Ryan J King, Kaymisha Knights, Ellen Koag, Nadezhda Kobko-Litskevitch, Giuseppe KoLoMonaco, Michael Kozhar, Nanami Kubota, Sheelta S Kumar, Lawrence Kwong, Rachel Kwong, Ingrid Lafontaine, Manolo Laiola, Isha Lamba, Hyunjung Lee, Lucy Lee, Yunmi Lee, Emily Leong, Marcus H Y Leung, Chenhao Li, Weijun Liang, Moses Lin, Yan Ling Wong, Priscilla Lisboa, Anna Litskevitch, Tracy Liu, Sonia Losim, Jennifer Lu, Simona Lysakova, Gustavo Adolfo Malca Salas, Denisse Maldonado, Krizzy Mallari, Tathiane M Malta, Maliha Mamun, Yuk Man Tang, Sonia Marinovic, Brunna Marques, Nicole Mathews, Yuri Matsuzaki, Madelyn May, Elias McComb, Adiell Melamed, Wayne Menary, Ambar Mendez, Katterinne N Mendez, Irene Meng, Ajay Menon, Mark Menor, Nancy Merino, Cem Meydan, Karishma Miah, Tanja Miketic, Eric Minwei Liu, Wilson Miranda, Athena Mitsios, Natasha Mohan, Mohammed Mohsin, Karobi Moitra, Laura Molina, Eftar Moniruzzaman, Sookwon Moon, Isabelle de Oliveira Moraes, Maritza S Mosella, Maritza S Mosella, Josef W Moser, Christopher Mozsary, Amanda L Muehlbauer, Oasima Muner, Muntaha Munia, Naimah Munim, Tatjana Mustac, Kaung Myat San, Areeg Naeem, Mayuko Nakagawa, Masaki Nasu, Bryan Nazario, Narasimha Rao Nedunuri, Aida Nesimi, Aida Nesimi, Gloria Nguyen, Hosna Noorzi, Avigdor Nosrati, Houtan Noushmehr, Diana N. Nunes, Kathryn O’Brien, Niamh B O’Hara, Gabriella Oken, Rantimi A Olawoyin, Kiara Olmeda, Itunu A Oluwadare, Tolulope Oluwadare, Jenessa Orpilla, Jacqueline Orrego, Melissa Ortega, Princess Osma, Israel O Osuolale, Oluwatosin M Osuolale, Rachid Ounit, Christos A Ouzounis, Subhamitra Pakrashi, Rachel Paras, Andrea Patrignani, Ante Peros, Sabrina Persaud, Anisia Peters, Robert A Petit III, Adam Phillips, Lisbeth Pineda, Alketa Plaku, Alma Plaku, Brianna Pompa-Hogan, Max Priestman, Bharath Prithiviraj, Sambhawa Priya, Phanthira Pugdeethosal, Benjamin Pulatov, Angelika Pupiec, Tao Qing, Saher Rahiel, Savlatjon Rahmatulloev, Kannan Rajendran, Aneisa Ramcharan, Adan Ramirez-Rojas, Shahryar Rana, Prashanthi Ratnanandan, Timothy D Read, Hugues Richard, Alexis Rivera, Michelle Rivera, Alessandro Robertiello, Courtney Robinson, Anyelic Rosario, Kaitlan Russell, Timothy Ryan Donahoe, Krista Ryon, Thais S Sabedot, Thais S Sabedot, Mahfuza Sabina, Cecilia Salazar, Jorge Sanchez, Ryan Sankar, Paulo Thiago de Souza Santos, Zulena Saravi, Thomas Saw Aung, Thomas Saw Aung, Nowshin Sayara, Steffen Schaaf, Anna-Lena M Schinke, Ralph Schlapbach, Jason R Schriml, Felipe Segato, Marianna S. Serpa, Heba Shaaban, Maheen Shakil, Hyenah Shim, Yuh Shiwa, Shaleni K Singh, Eunice So, Camila Souza, Jason Sperry, Kiyoshi Suganuma, Hamood Suliman, Jill Sullivan, Jill Sullivan, Fumie Takahara, Isabella K Takenaka, Anyi Tang, Emilio Tarcitano, Mahdi Taye, Alexis Terrero, Andrew M Thomas, Sade Thomas, Masaru Tomita, Xinzhao Tong, Jennifer M Tran, Catalina Truong, Stefan I Tsonev, Kazutoshi Tsuda, Michelle Tuz, Carmen Urgiles, Brandon Valentine, Hitler Francois Vasquez Arevalo, Valeria Ventorino, Patricia Vera-Wolf, Sierra Vincent, Renee Vivancos- Koopman, Andrew Wan, Cindy Wang, Samuel Weekes, Xiao Wen Cai, Johannes Werner, David Westfall, Lothar H Wieler, Michelle Williams, Silver A Wolf, Brian Wong, Tyler Wong, Hyun Woo Joo, Rasheena Wright, Ryota Yamanaka, Jingcheng Yang, Hirokazu Yano, George C Yeh, Tsoi Ying Lai, Laraib Zafar, Amy Zhang, Shu Zhang, Yang Zhang, Yuanting Zheng,

## 9 Supplemental Materials

### 9.1 Supplemental Methods

#### 9.1.1 DNA Extraction from Isohelix swabs using ZymoBiomics 96 MagBead

The Isohelix swab head and the entire 400 *μ*l of DNA/RNA Shield-solubilized sample were transferred into ZR BashingBead Lysis Tubes (0.1 & 0.5 mm) (Cat# S6012-50) to which an additional 600 *μ*l of DNA/RNA Shield was added. Mechanical lysis using bead beating was performed on a maximum of 18 samples simultaneously using the Scientific Industries Vortex-Genie 2 with Horizontal-(24) Microtube Adapter (Cat # SI-0236 and SI-H524) at maximum power for 40 minutes. The resulting lysate (400 *μ*l) was transferred to Nunc™ 96-Well Polypropylene DeepWell Storage Plates (Cat # 278743), followed by DNA extraction using the ZymoBIOMICS 96 MagBead Kit (Lysis Tubes) (Catalog # D4308) on the Hamilton Star according to manufacturer instructions.

#### 9.1.2 DNA extraction from Copan swabs using MoBio PowerSoil®DNA

Droplets in the Copan Liquid Amies Elution Swab tube (ESwab, Copan Diagnostics, Cat.:480C (http://goo.gl/8a9uCP)) were spun down at 300rpm/1min. Next, the swab pad was transferred to a Mo- Bio PowerSoil®DNA vial containing beads using sterile scissors, which we sterilized by flaming with 100% ethanol. The remaining 400-500*μ*l Copan Amines liquid has been transferred into an Eppendorf tube and centrifuged at full speed to collect bacteria and debris in a pellet. The pellet was finally transferred to the same MoBio PowerSoil®DNA vial also containing the corresponding swab pad. MoBio PowerSoil®DNA Isolation Kit, Cat.:12888-100 (https://www.qiagen.com/us/resources/resourcedetail?id=5c00f8e4-c9f5-4544-94fa-653a5b2a6373&lang=en) was used according to manufacturer’s instructions except for the following modifications:

Both swab and pellet have been re-suspended with 135*μ*l C1 buffer (MoBio PowerSoil®DNA). Sample homogenization was performed using either TissueLyser II (Qiagen) with 2 cycles of 3 minutes at 30Hz (https://goo.gl/hBg8Lb), or using the Vortex-Genie 2 (Vortex Catalog #13000-V1-24) adaptor and vortex at maximum speed for 10 minutes. The sequencing centers in Stockholm and Shanghai used different procedures for homogenization. Stockholm used a method based on MPI FASTPREP, while Shanghai added 0.6 grams of 100-micron zirconium-silica beads to 2ml tubes containing the swab pad and the media, followed by bead beating for 1 min. The eluted samples have been additionally purified and concentrated by Beckmann Coulter Agencourt AMPure XP (Cat.:A63881) purification (1.8X) and eluted into 12*μ*l – 50*μ*l elution buffer. Subsequently, DNA was quantified using Qubit® dsDNA HS Assay (Catalog #Q32854).

#### 9.1.3 DNA extraction using Promega Maxwell

We added 300*μ*l Promega Maxwell Lysis buffer and 30*μ*l Promega Maxwell Proteinase K to Copan swab heads or Isohelix swab heads and transferred the swabs back to their respective collection tube. For lysis the sample tubes containing the swabs and the lysis mixture were incubated in a water bath at 54C for 30min. Following lysis, Copan swab heads were cut off their stem using sterile scissors and transferred into a filter tube (Promega V4745). The filter containing the swab was placed into a 2ml Eppendorf tube and spun down at full speed for 2min. This step is necessary since the Copan swab material consists of a foam, which harbors the main liquid containing the extracted DNA. Next, the eluate has been combined with the corresponding sample tube media and added to the first well of the cartridge (Maxwell® RSC Buccal Swab kit AS1640). Cartridges were processed using the Maxwell® RSC Instrument (AS4500) following the manufacturer’s default instructions. Extracted DNA was eluted in 50*μ*l Promega Elution Buffer and stored at −80C.

The matrix tubes containing the Isohelix swabs and the lysis buffer have been vortexed at full speed for one minute. The Isohelix swab head material is a non-porous material, which allows for easy collection of the lysate. We transferred the lysate to the first cartridge of the Maxwell® RSC Blood DNA KitAS1400 using syringes (BD 3 mL Syringes with 18G x 1.5” Luer Lok Tip Blunt Fill Needles) and ran the Promega Maxwell using the Blood program according to manufacturer’s instructions. Samples were subsequently eluted in 50*μ*l elution buffer and stored at −80C.

**Table S1:**
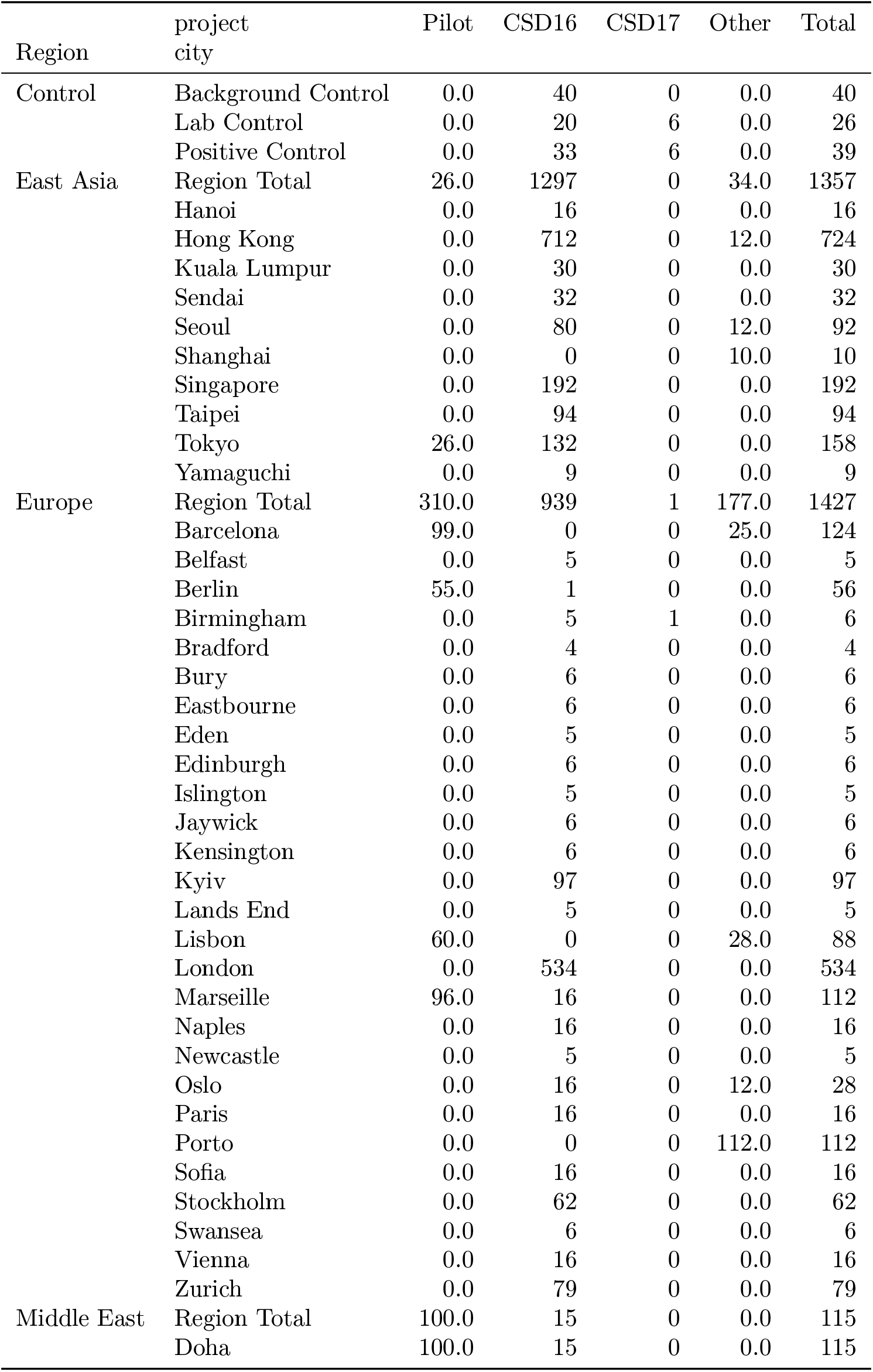
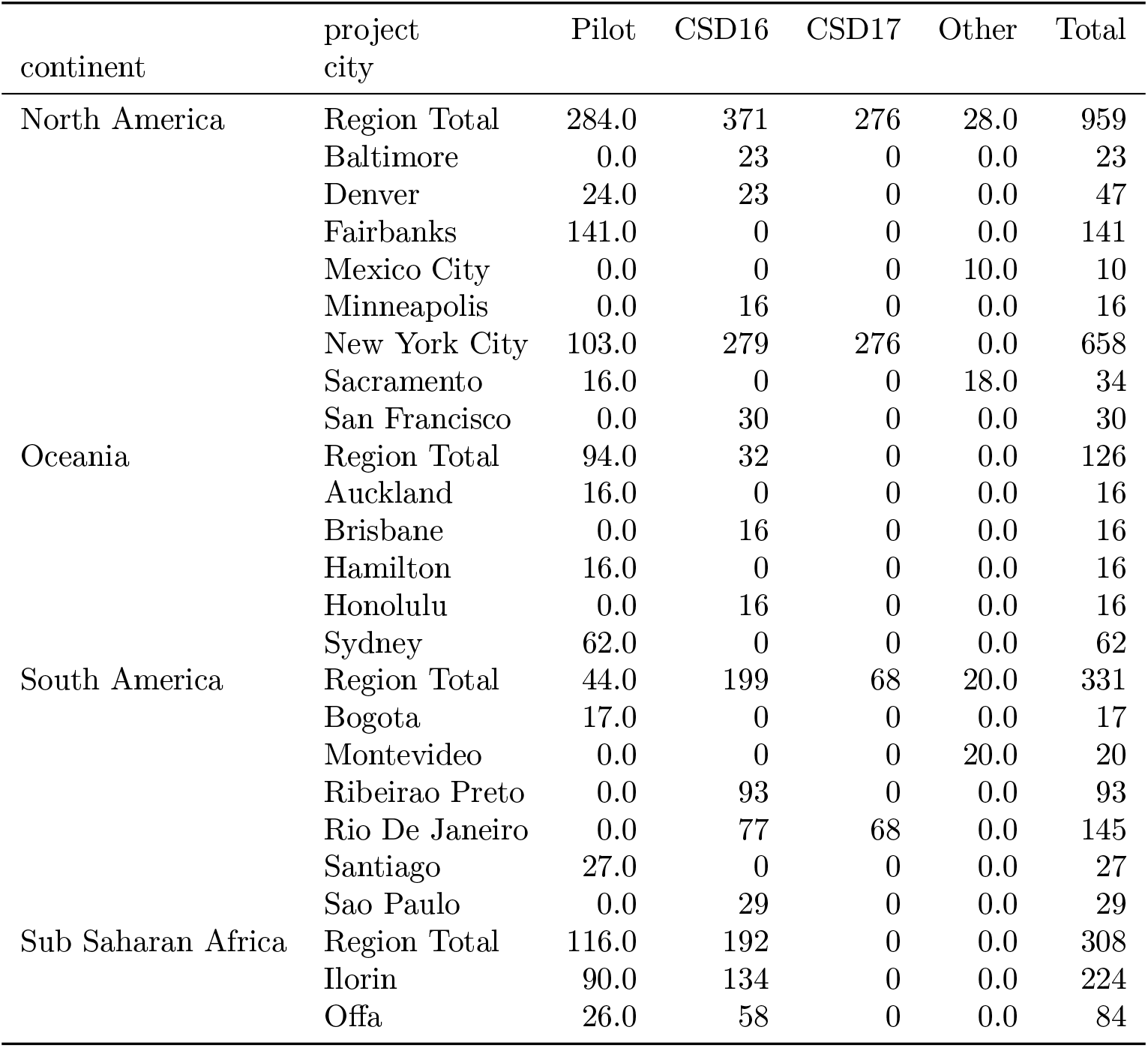
Sample Counts

**Table S2:**
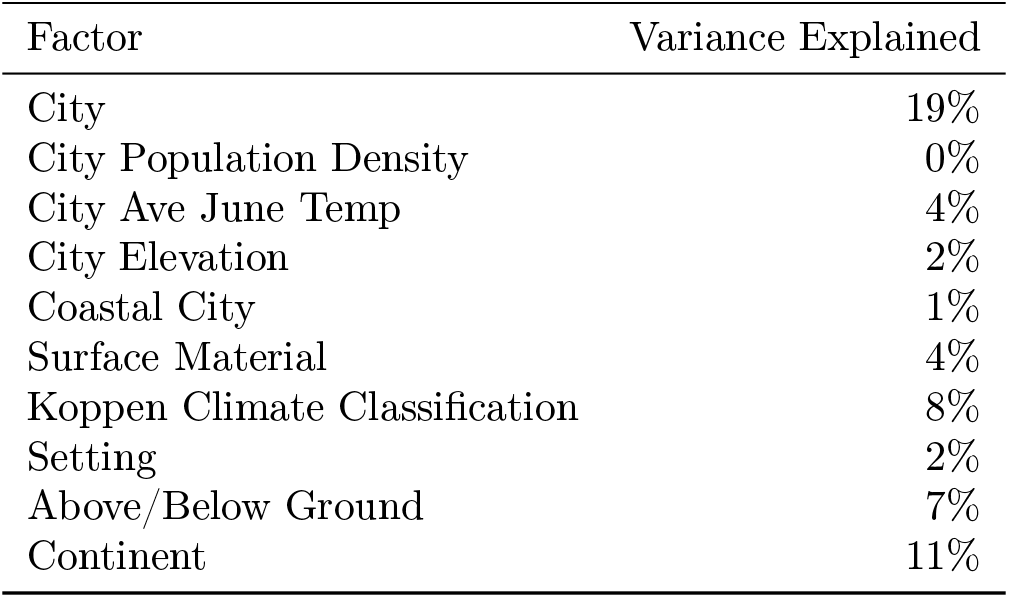
Covariate Variance. The sample variance that can be explained by each factor, in isolation.

**Figure S1:**
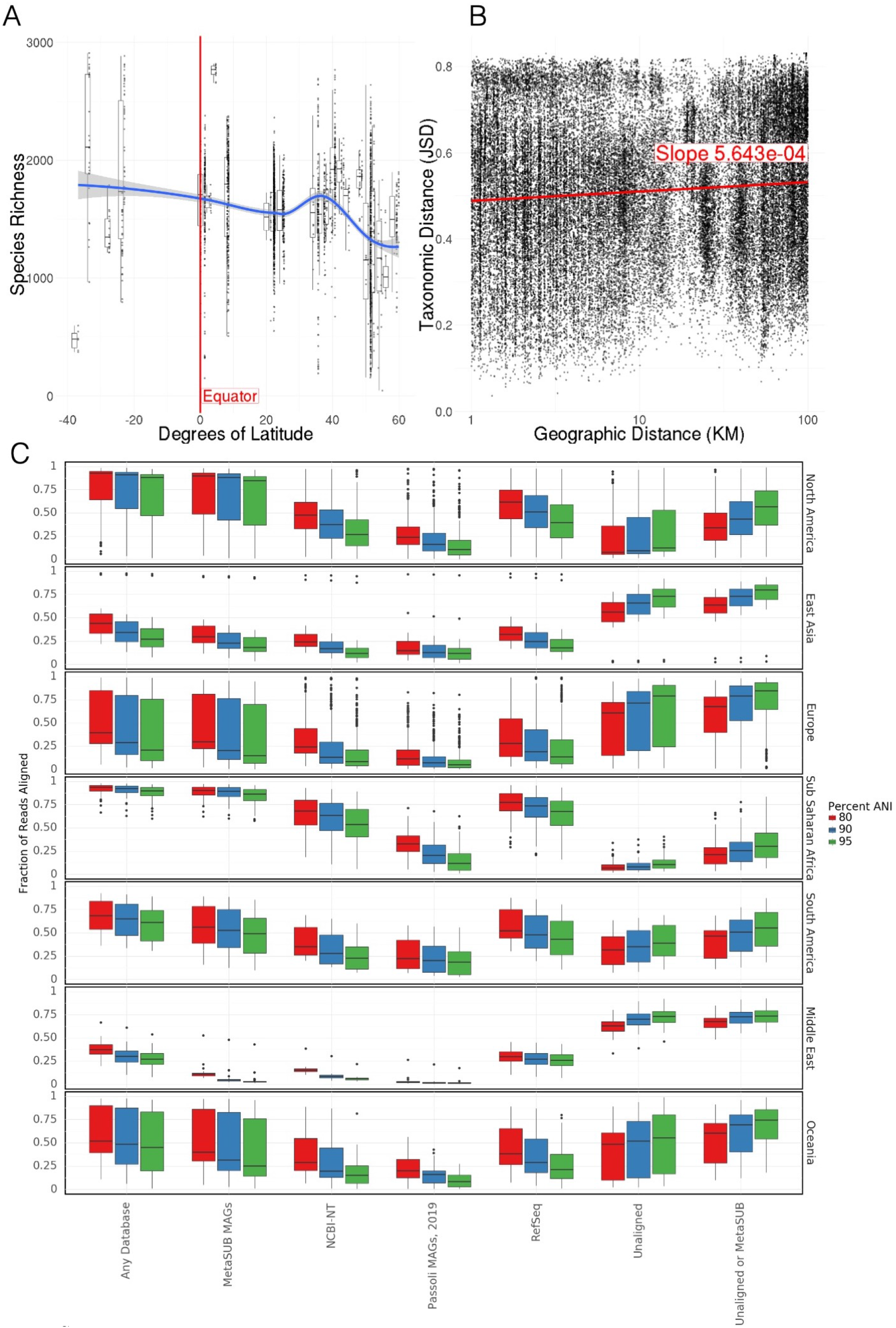
Ecological relationships with taxa. A) Correlation between species richness and latitude. Richness decreases significantly with latitude B) Neighbourhood effect. Taxonomic distance weakly correlates with geographic distance within cities. C) Fraction of reads assigned to different databases by BLAST for each region, at different levels of average nucleotide identity

**Figure S2:**
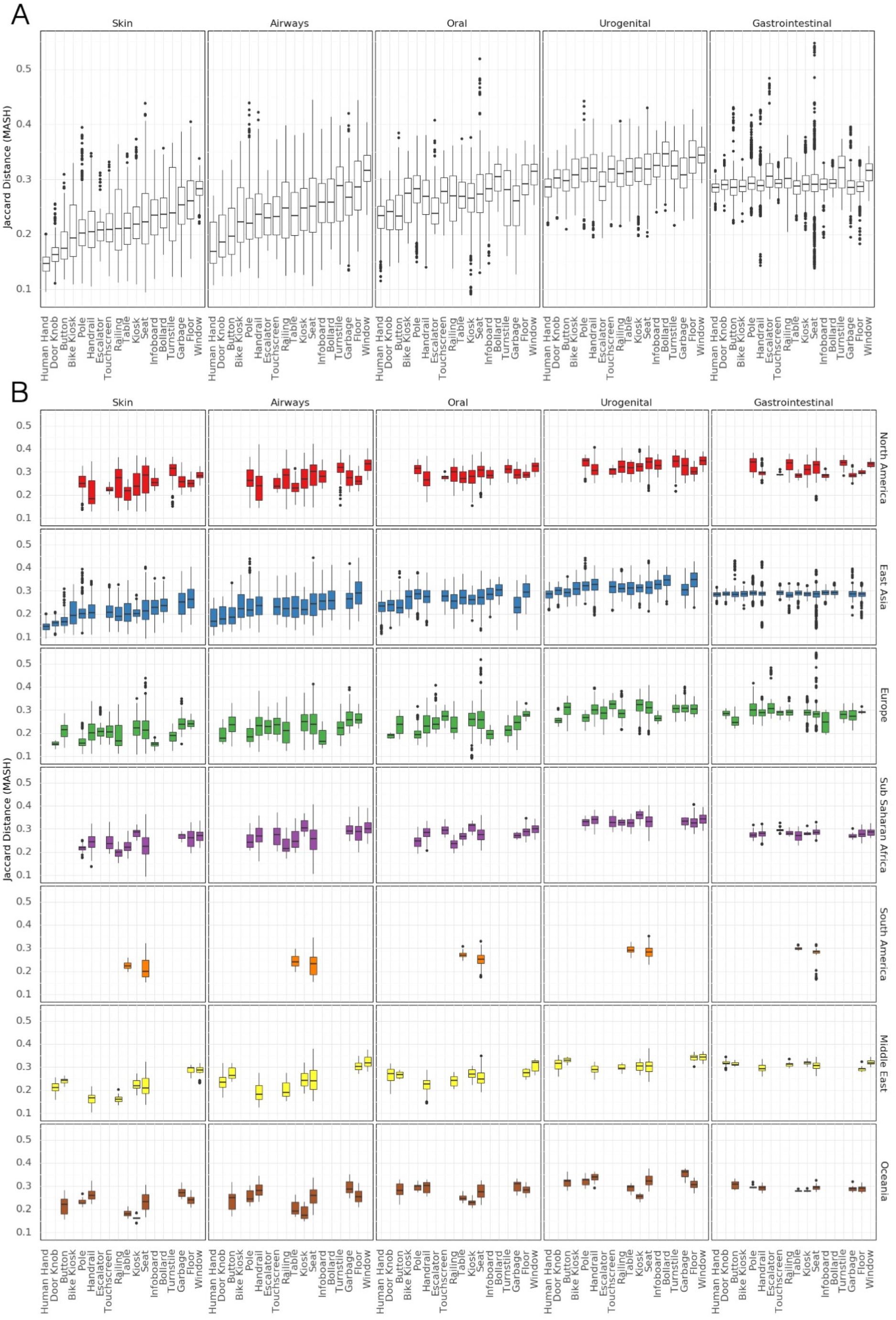
Comparison to Human Microbiome Project. A) Jaccard similarity of MASH indices to HMP samples for different surface types. B) Jaccard similarity of MASH indices to HMP samples for different surface types by region.

**Figure S3:**
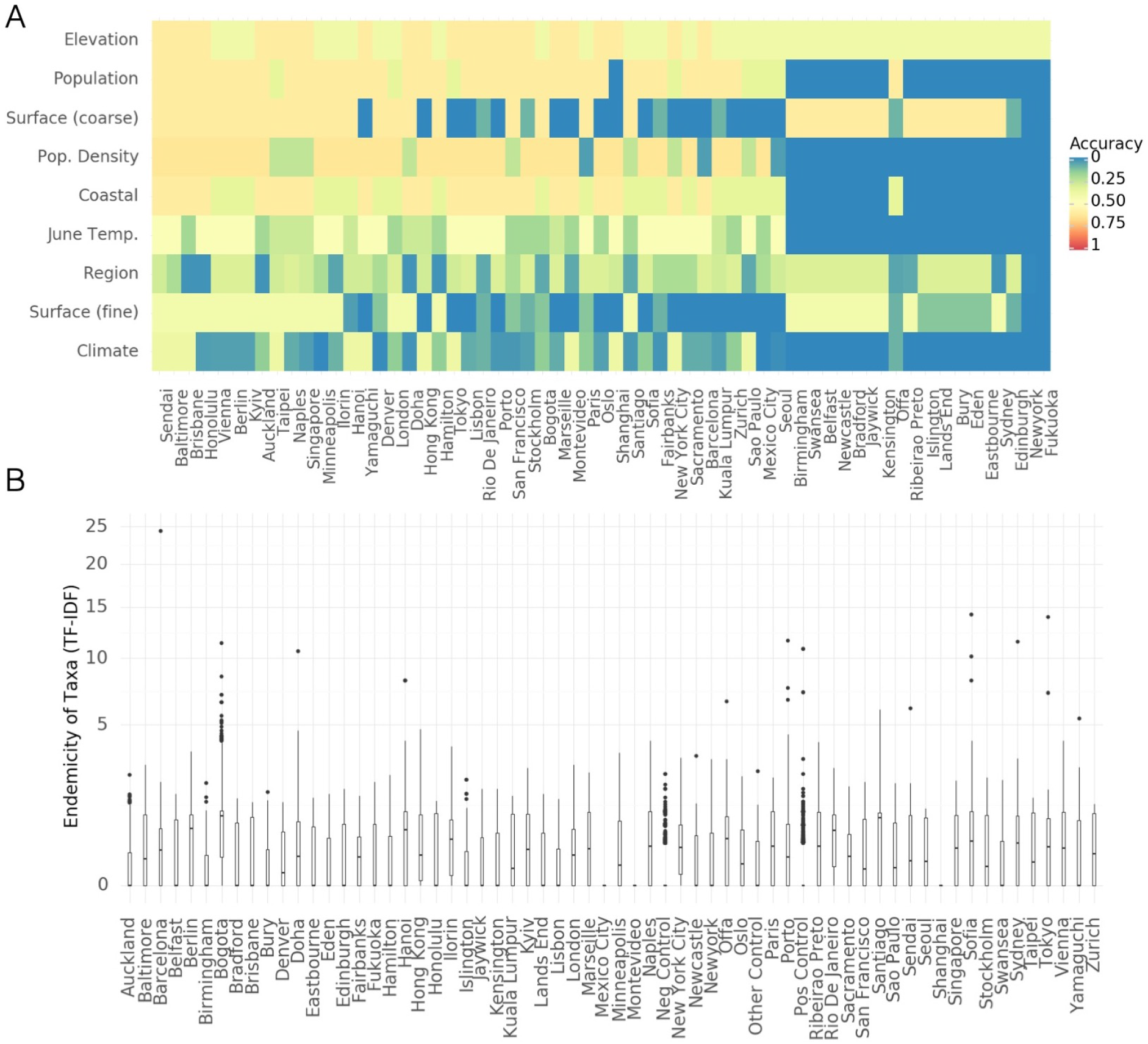
Microbial Signatures, supplemental. A) Classification accuracy that would be achieved by a random model predicting features (rows) for held out cities (columns) B) Endemicity Score (Term Frequency Inverse Document Frequency for taxa in cities

**Figure S4:**
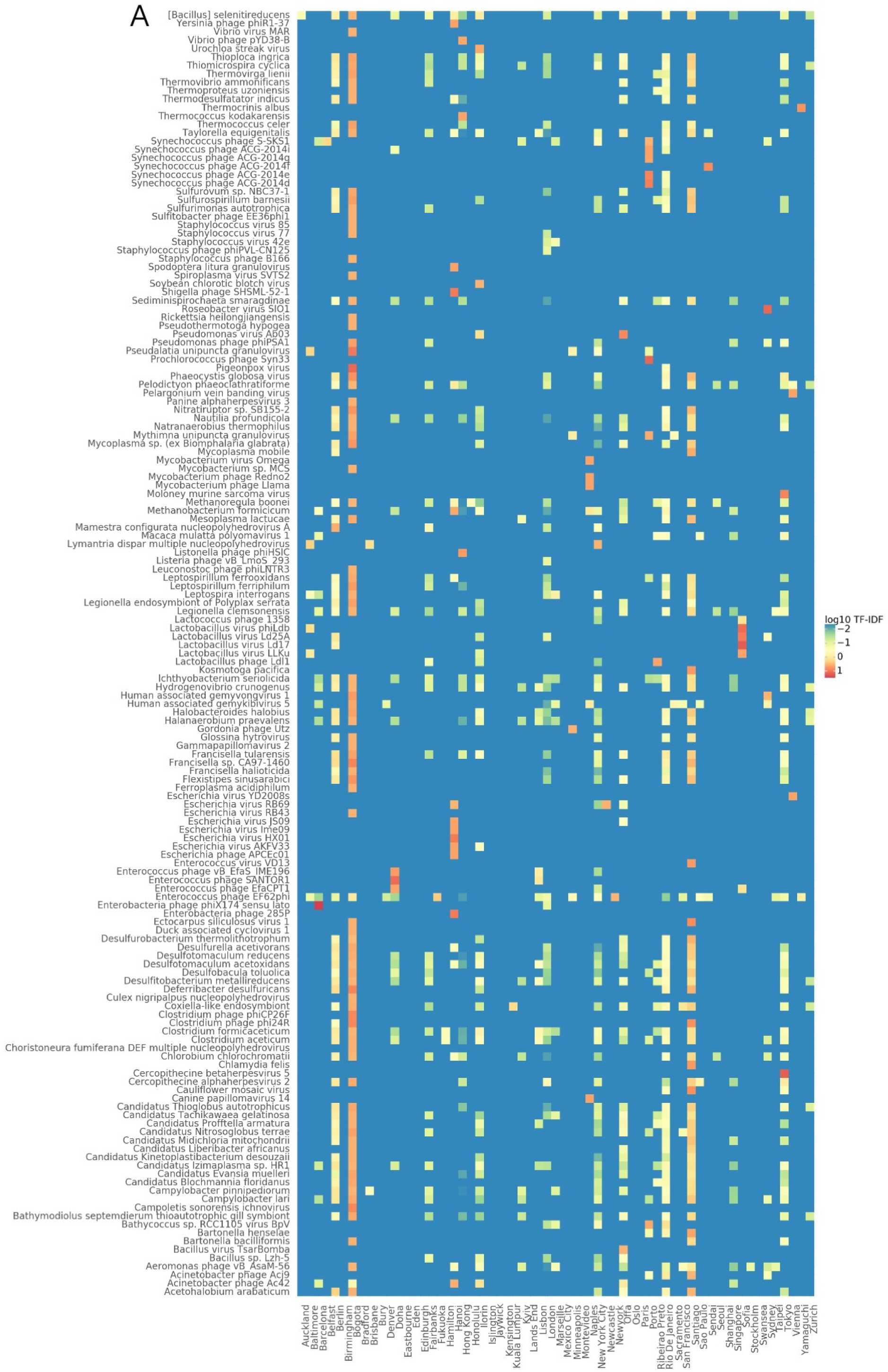
Endemicity scores of particular taxa. A) Heatmap showing the endemicity scores (term-frequency inverse document frequency) for taxa in different cities. This table is filtered to show only taxa with high endemicity scores in at least one city.

**Figure S5:**
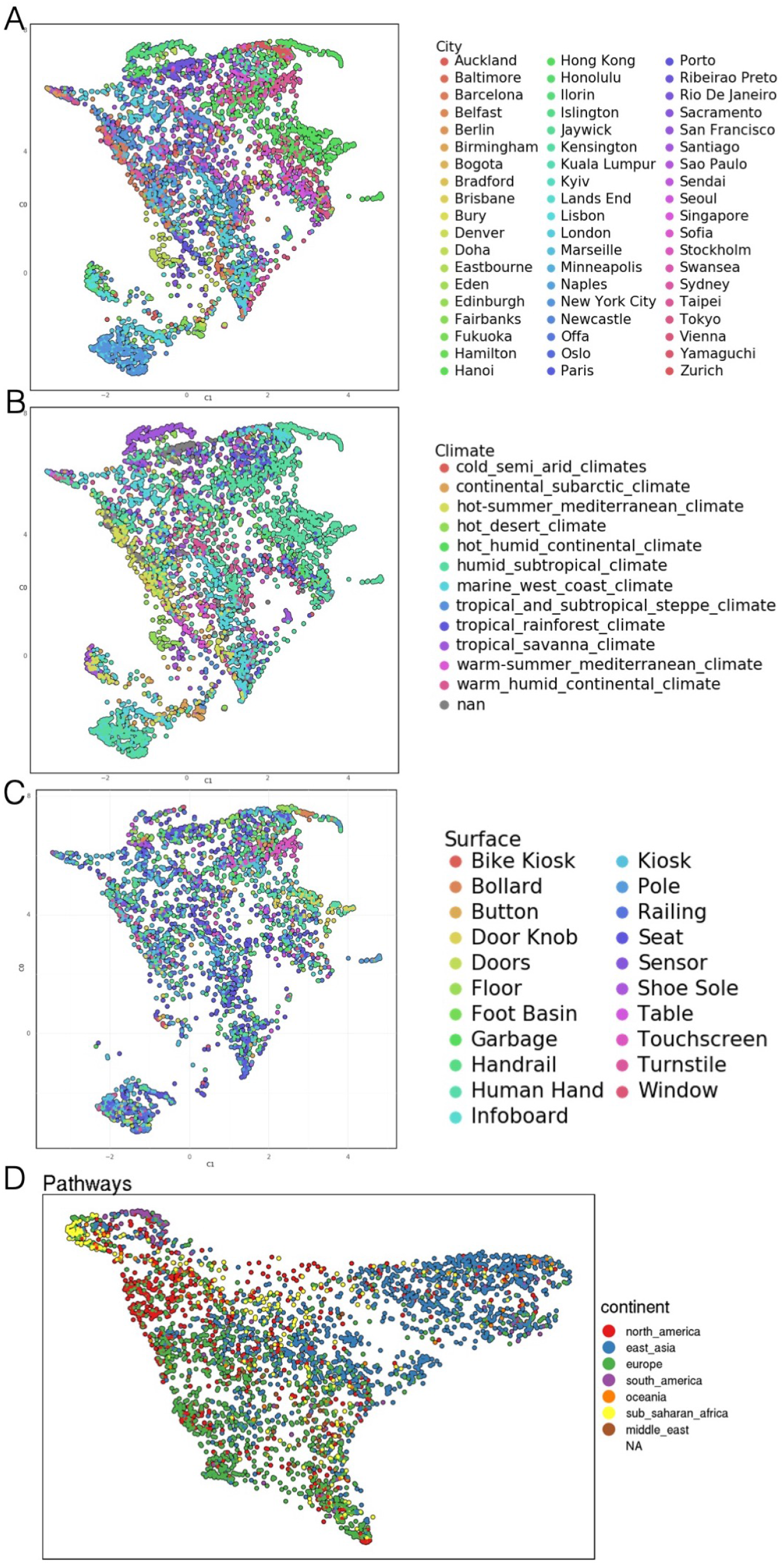
A) UMAP of taxonomic profiles colored by city B) UMAP of taxonomic profiles colored by climate classification C) UMAP of taxonomic profiles colored by surface type D) UMAP of functional profiles colored by region

**Figure S6:**
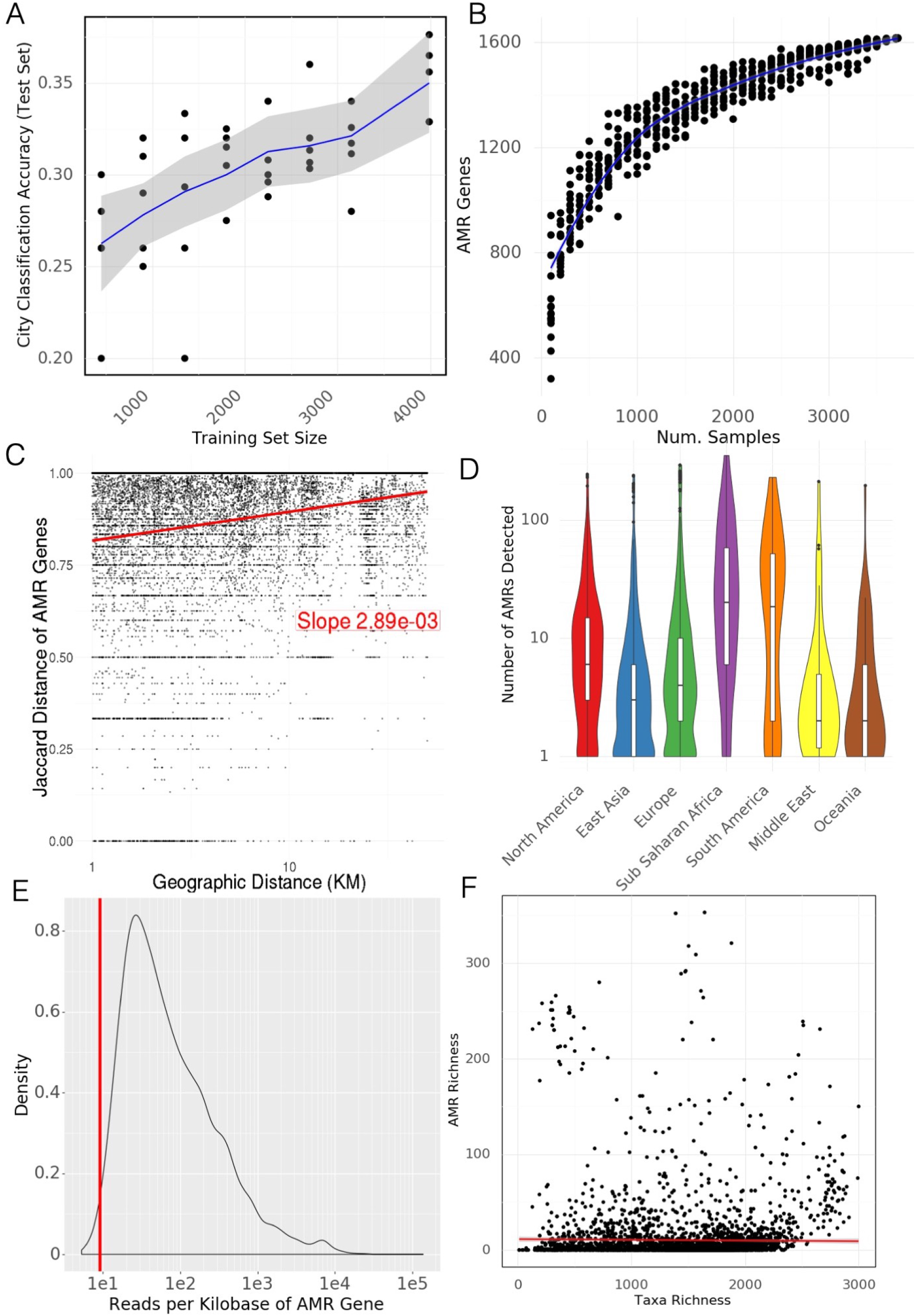
Antimicrobial Resistance Genes, supplemental. A) Classification accuracy of a random forest model predicting city labels for held out samples from antimicrobial resistance genes. B) Rarefaction analysis of antimicrobial resistance genes. Curve does not flatten suggesting we would identify more AMR genes with more samples. C) Neighbourhood effect. Jaccard distance of AMR genes weakly correlates with geographic distance within cities. D) Number of AMR genes detected for samples in each region. E) Distribution of reads per gene (normalized by kilobases of gene length) for AMR gene calls. The vertical red line indicates that 99% of AMR genes have more than 9.06 reads per kilobase and would still be called at a lower read depth.

**Figure S7:**
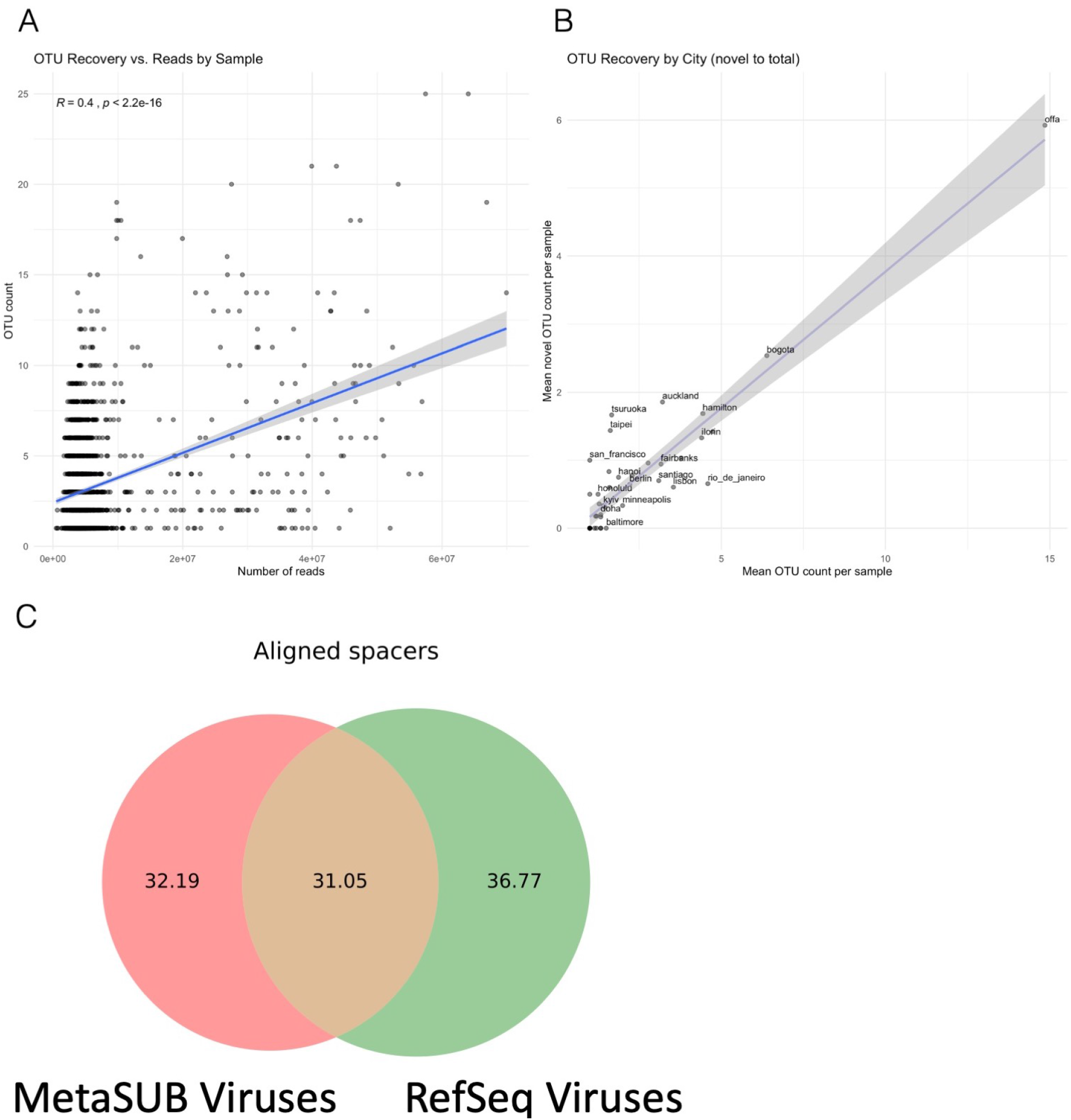
Novel biology, supplemental. A) Relation of read depth to the number of identified bacterial Metagenome Assembled Genomes (MAGs) in a sample. B) Discovery rate for baterial MAGs in each city. C) Total fraction of CRISPR spacers aligned to MetaSUB viral MAGs and viral genomes in RefSeq.

**Figure S8:**
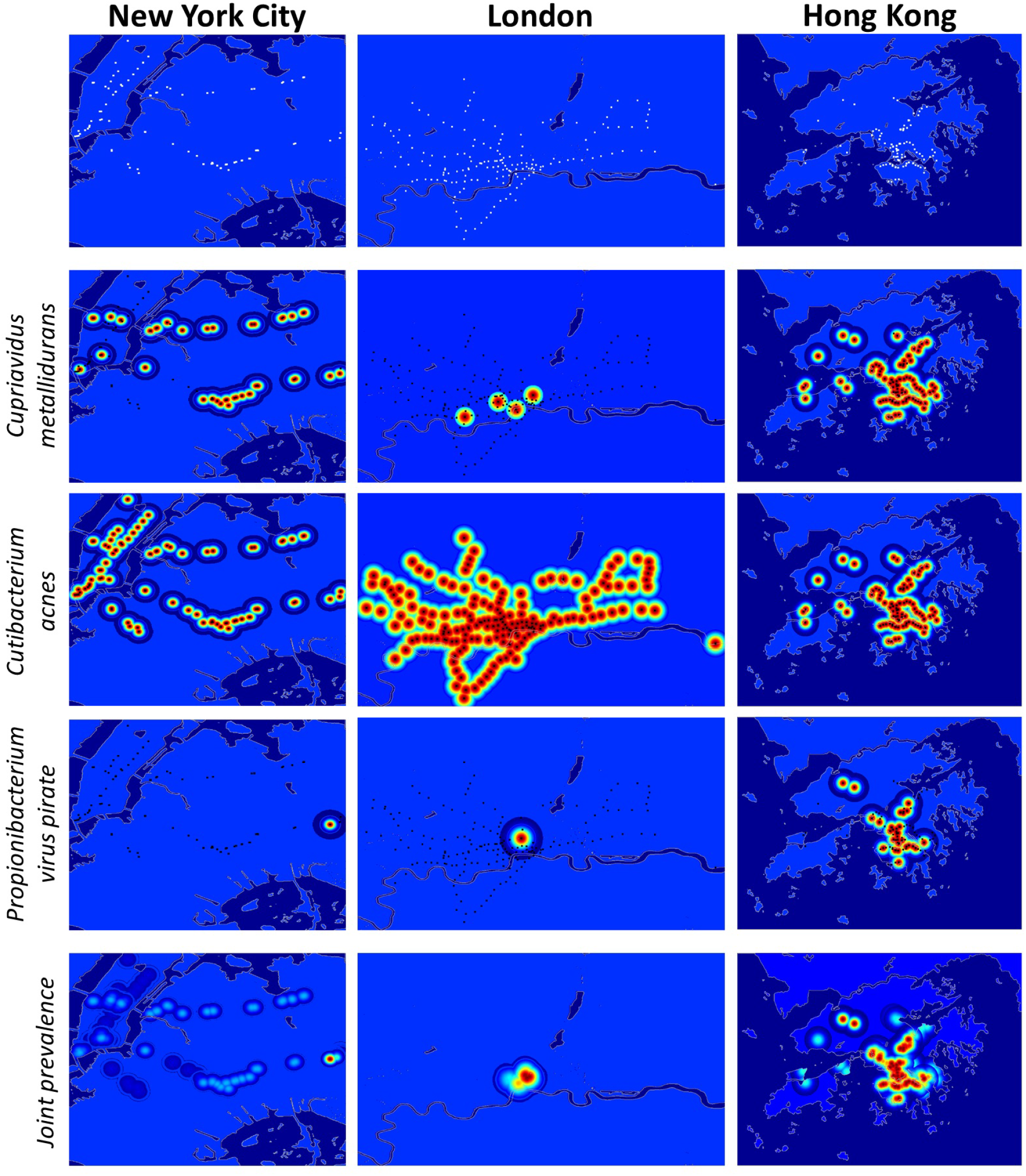
Example Geographic taxonomic Distributions. Distributions of taxa were estimated by fitting Gaussian distributions to sampling locations where the taxa was found with standard deviations based on the geographic distance between observations. Top Row) Sampling sites in three major cities Rows 2-4) Estimated distribution of different example species in major cities Row 5) Estimated distribution of three species together in major cities

**Figure S9:**
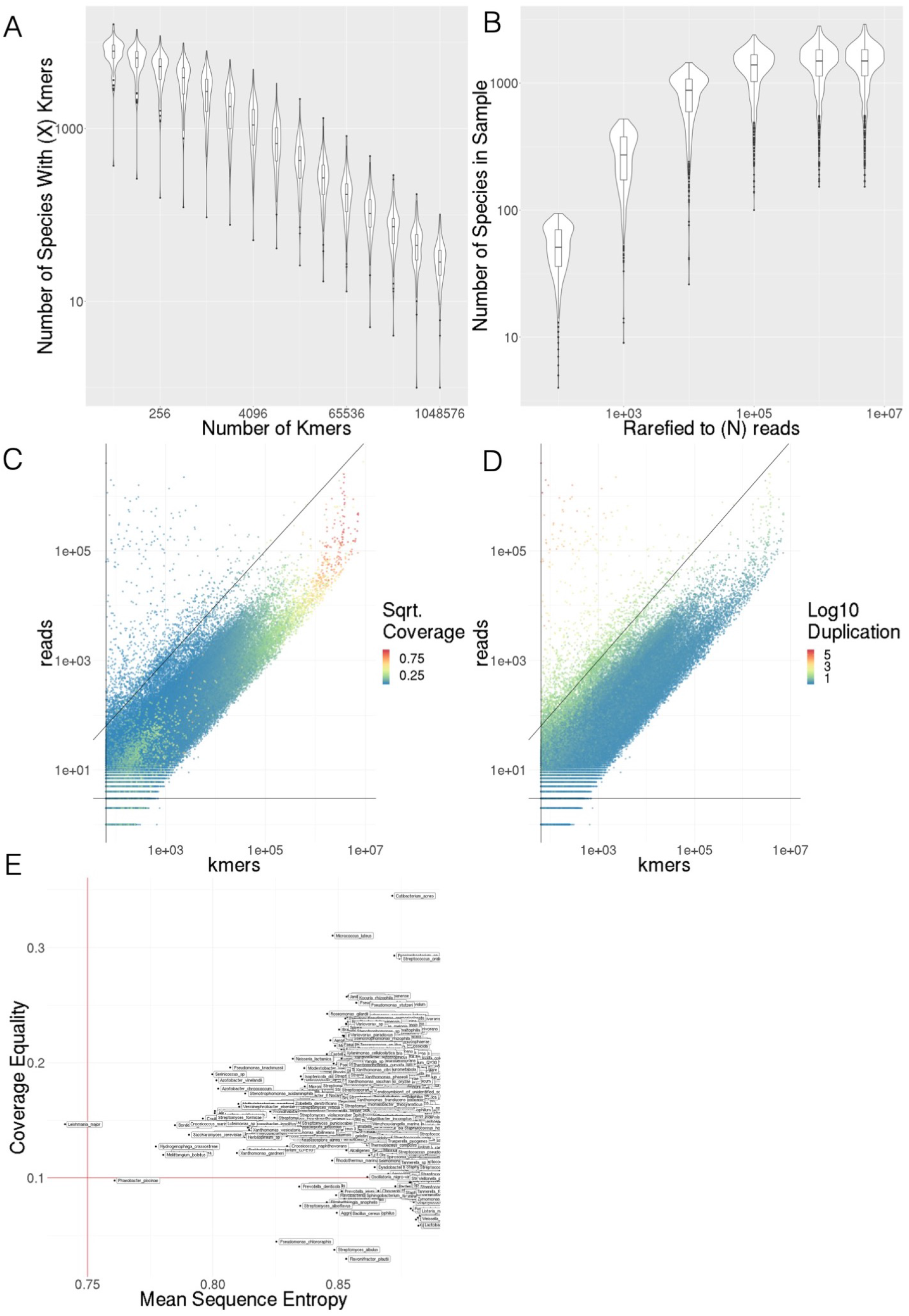
A) Number of species detected as *k*-mer threshold increases for 100 randomly selected samples B) Number of species detected as number of sub-sampled reads increase C) *k*-mer counts compared to number of reads for species level annotations in 100 randomly selected samples, colored by coverage of marker *k*-mer set D) *k*-mer counts compared to number of reads for species level annotations in 100 randomly selected samples, colored by average duplication of *k*-mers E) Comparison of Mean Sequence Entropy and Coverage Equality for core and sub-core taxa. Thresholds are shown by red lines.

**Figure S10:**
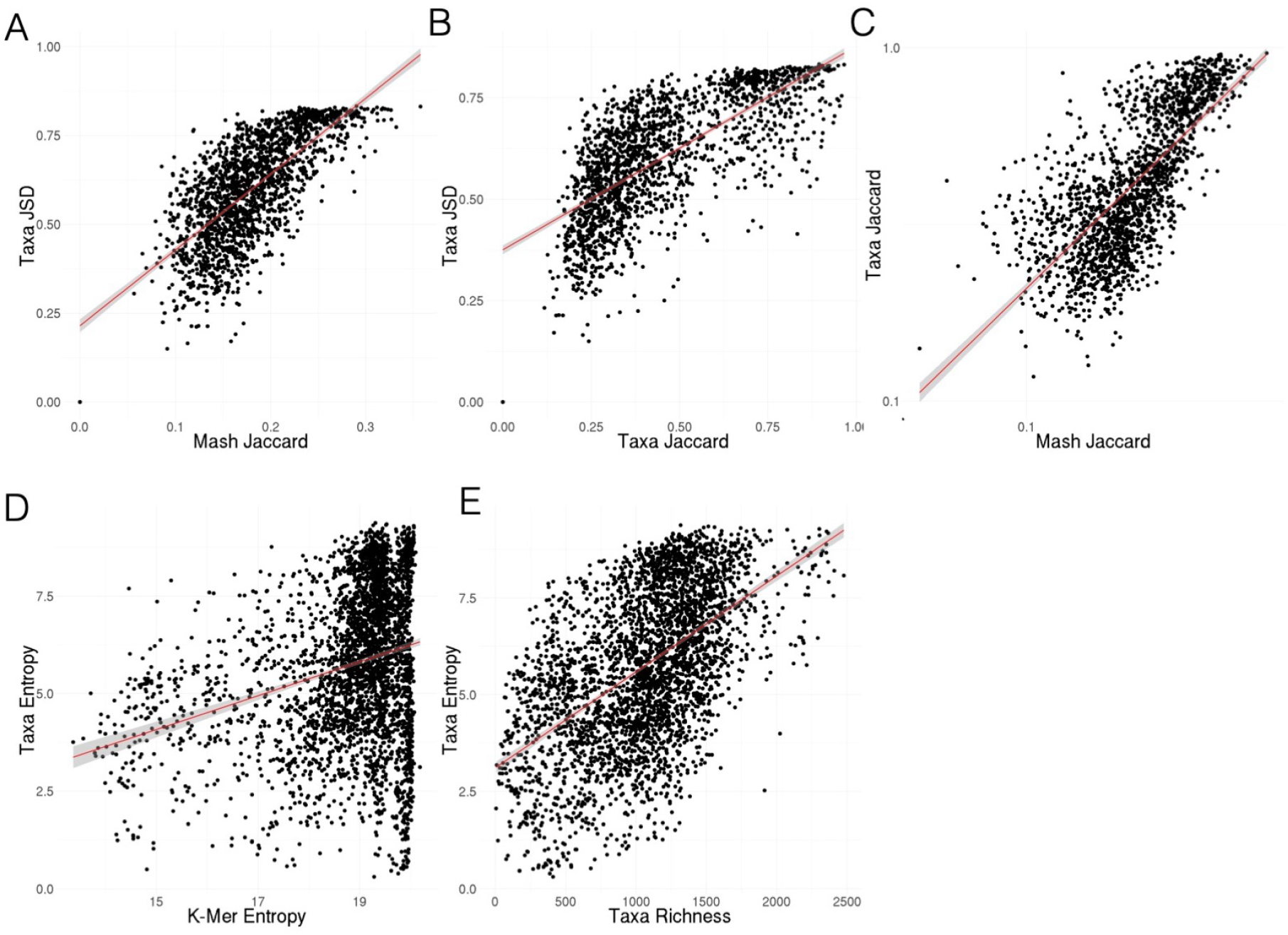
A) Jensen-Shannon Divergence of taxonomic profiles vs MASH Jaccard distance of *k*-mers B) Jensen-Shannon Divergence of taxonomic profiles vs Jaccard distance of taxonomic profiles. C) Jaccard distance of taxonomic profilesvs MASH Jaccard distance of *k*-mers D) Shannon’s Entropy of taxonomic profiles vs Shannon’s Entropy of *k*-mers E) Taxonomic richness (number of species) vs Shannon’s Entropy of taxonomic profiles

**Figure S11:**
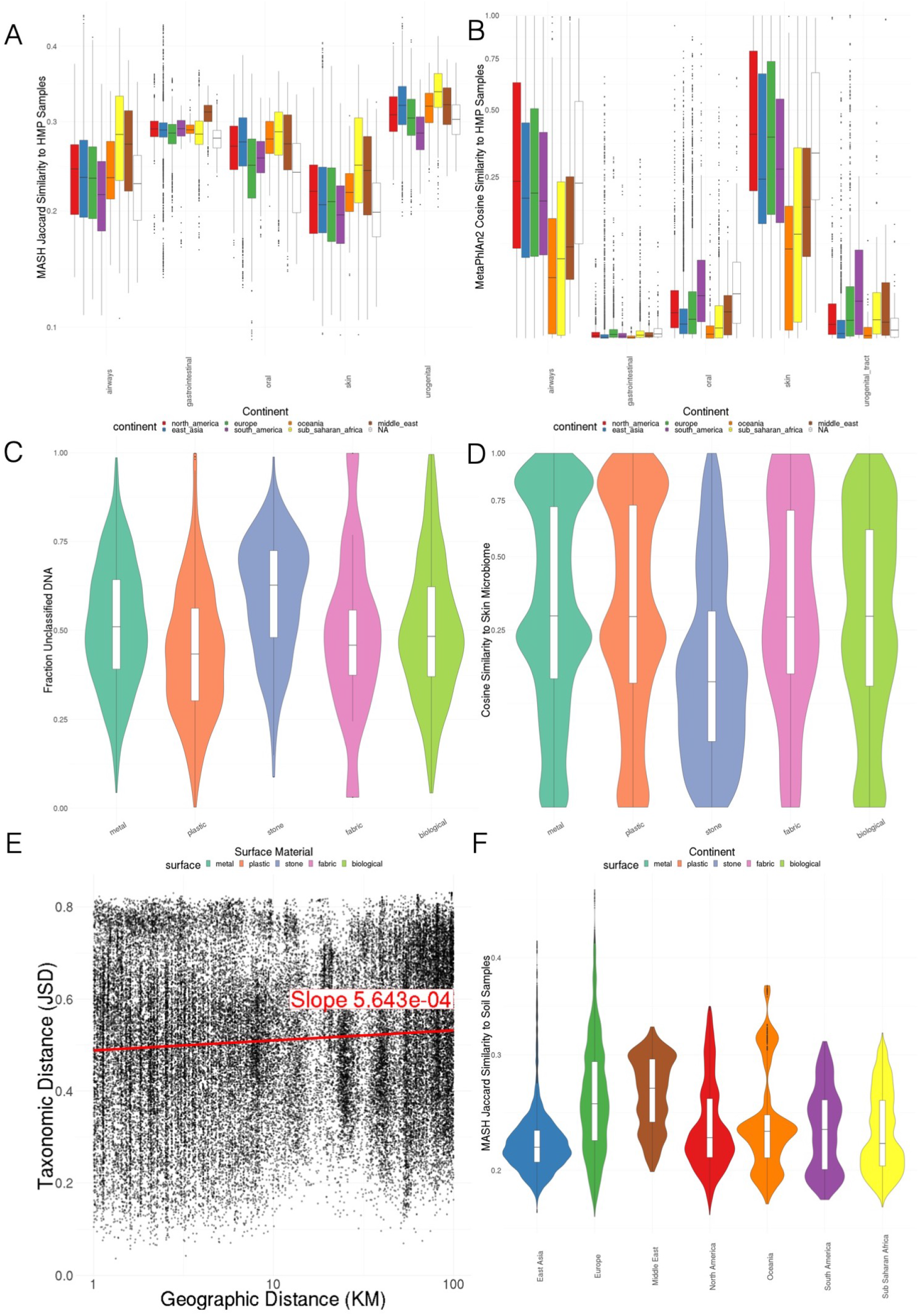
A) MASH *k*-mer Jaccard similarity to representative HMP samples, colored by continent B) MetaPhlAn v2.0 cosine similarity to representative HMP samples, colored by continent C) Fraction unclassified DNA by surface material D) Cosine similarity to MetaPhlAn v2.0 skin microbiome profile by surface E) Jensen-Shannon distance between pairs of taxonomic profiles vs Geographic Distance F) MASH *k*-mer Jaccard similarity to representative soil samples, colored by continent

**Figure S12:**
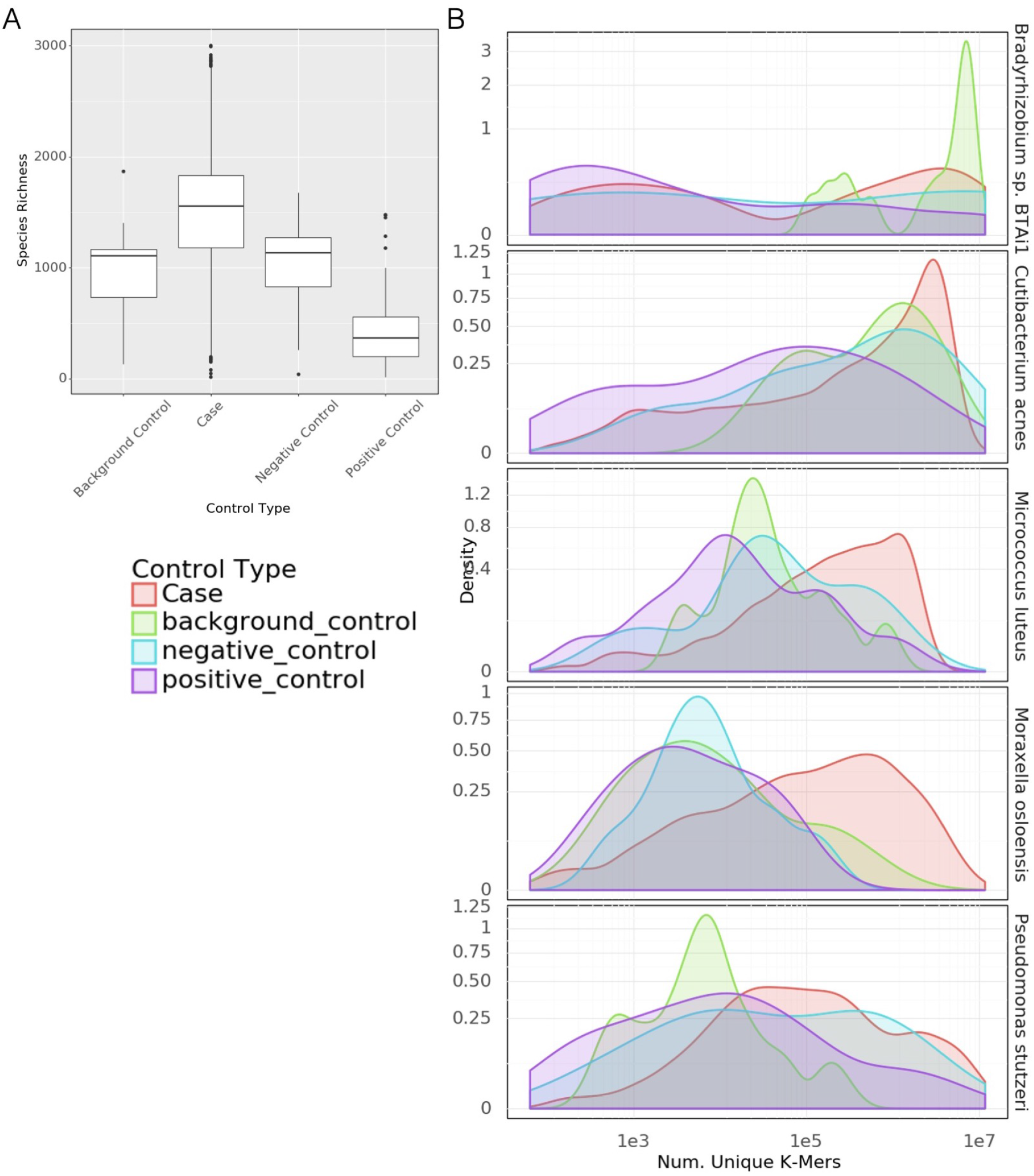
A) Taxonomic Richness in Cases vs. Types of Controls B) Distributions of *k*-mer counts in control types vs cases for 5 most abundant taxa. *k*-mer count is a marker of assignment confidence.

**Figure S13:**
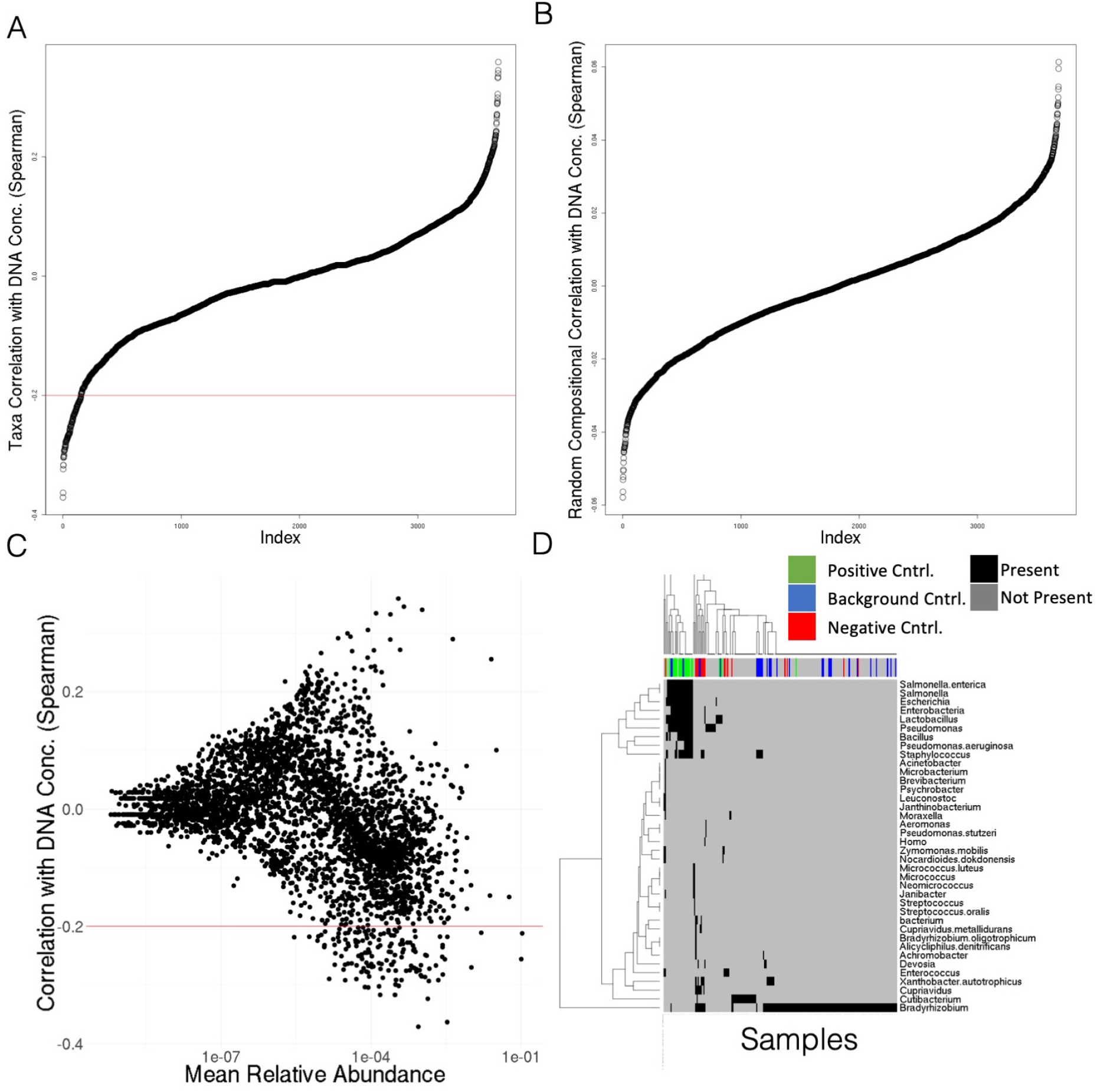
A) Correlation of taxonomic (species) relative abundances with DNA concentration B) Correlation of randomly generated compositional vectors with DNA concentration. Note the same shape but lower magnitude C) Correlation of taxa with DNA Concentration vs the mean relative abundance of that taxa D) Presence (black) absence (grey) heatmap of taxa found in controls and other samples. Colored bar at top, red are negative controls, blue are background, green are positive. Case samples with homology are grey. Case samples without homology to control sequences are not shown.

**Figure S14:**
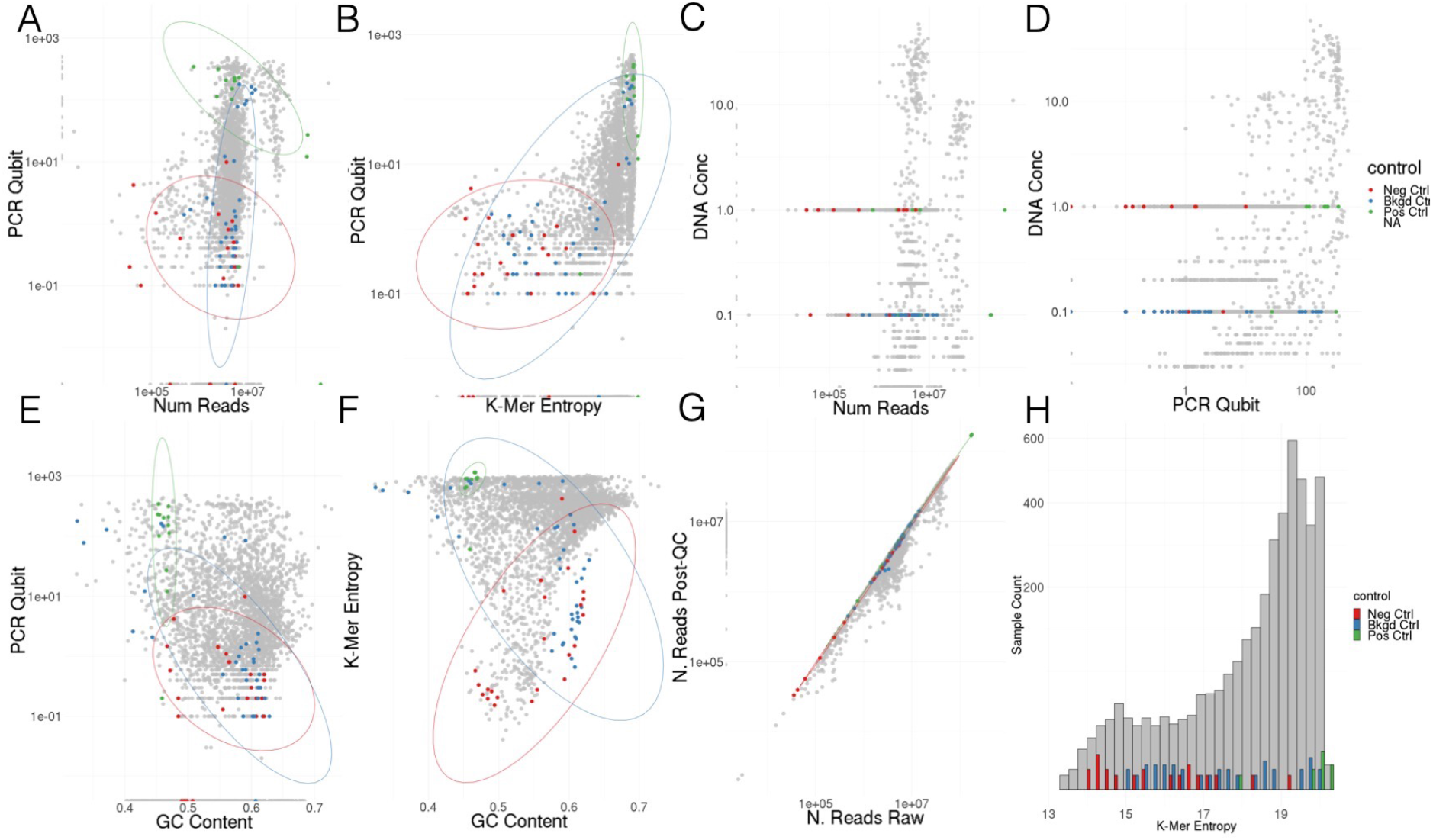
Comparisons of different seuqeuncing quality control metrics with controls marked. A-F) Comparisons of the raw reads, PCR Qubit scores, manually recorded DNA concentrations, *k*-mer Shannon entropy, and GC fraction of quality controlled reads G) Comparison of read counts before and after quality control but before human reads were removed H) Histogram showing the number of samples with different *k*-mer entropies.

**Figure S15:**
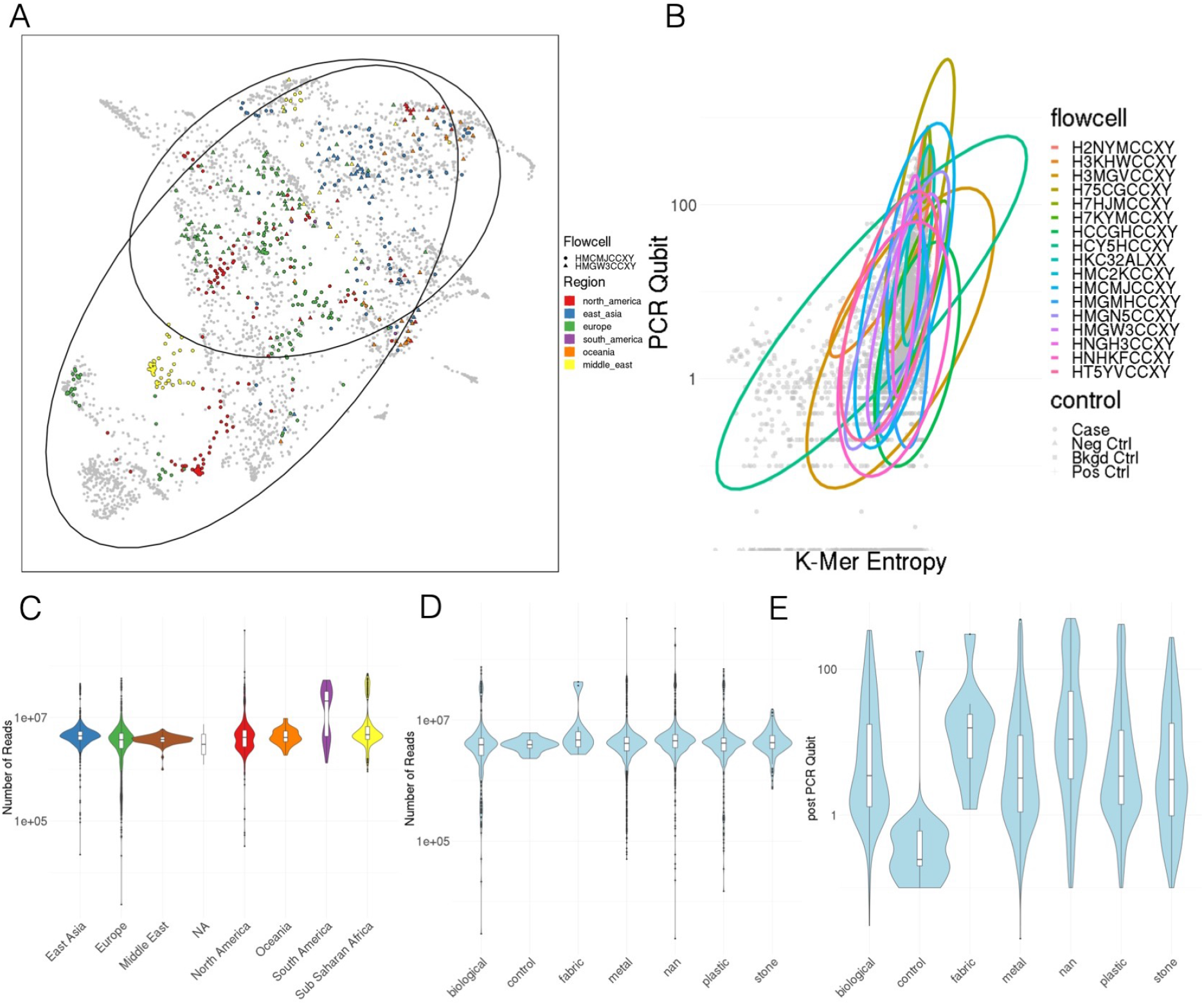
A) UMAP of taxonomic profiles from geographically diverse flowcells B) Flowcells vs quality control metrics C) Number of reads by region D) number of reads by surface material E) PCR Qubit by surface material

